# Cell Size Modulates SBF and Whi5 Chromatin Binding to Regulate the *Start* of the Budding Yeast Cell Cycle

**DOI:** 10.64898/2026.02.15.705999

**Authors:** H. Zheng, A. P. Gutierrez Alejandre, M. Shafieidarabi, M. P. Swaffer, Z. W. El-Hajj, M. Vera, J.M. Skotheim, R Reyes-Lamothe

**Affiliations:** Biology Department, McGill University, Montreal, Qc, Canada; Department of Biology, Stanford University, Stanford, CA, USA; Wellcome Centre for Cell Biology, University of Edinburgh, Edinburgh, UK; Biochemistry Department, McGill University, Montreal, Qc, Canada; Chan Zuckerberg Biohub, San Francisco, CA 94158, USA; Microbiology & Immunology Department, McGill University, Montreal, Qc, Canada; Centre de Recherche en Biologie Structurale, McGill University, Montreal, Qc, H3G 0B1, Canada

## Abstract

Cell growth and division are tightly coordinated to cell size. In budding yeast, increasing cell size promotes the G1/S transition, called *Start*, by activating the transcription factor SBF, which drives a large fraction of cell-cycle–dependent gene expression. Part of this regulation arises because the concentration of the SBF inhibitor Whi5 decreases as cells grow. However, cells lacking Whi5 can still maintain a relatively accurate size when the SBF activator Cln3 is also removed, indicating that there are additional size control mechanisms. To understand how cell size is mechanistically translated into the activity of SBF-regulated promoters, we quantified the binding kinetics of Whi5 and SBF in live cells using single-molecule fluorescence microscopy. We found that increasing cell size is associated with both a decreased chromatin affinity of Whi5 and an increased chromatin affinity of SBF, accompanied by a higher SBF:Whi5 cell copy-number ratio. Chromatin-binding trends under basal and Whi5 overexpression conditions indicate that Whi5 restricts SBF association with chromatin. The transition point at which SBF binding overtakes Whi5 binding coincides with the onset of the expression of the G1 cyclins *CLN1* and *CLN2*, two SBF targets that are important for committing cells to division. Reduced Whi5 binding reflects changes in its chromatin-association rate, as Whi5 and SBF dwell times on chromatin remain ∼10 s and are largely independent of cell size. Together, these results show how changes in SBF and Whi5 abundance and chromatin association transmit cell size information to the genome to regulate the size-dependent *Start* transition in budding yeast.

## INTRODUCTION

Cell size homeostasis is a universal feature of biology^1,2^. While cell size homeostasis is commonly observed, the underlying molecular mechanisms have been difficult to isolate and determine. The understanding of molecular mechanisms is most developed in the budding and fission yeast model organisms^3–6^. In the budding yeast, *Saccharomyces cerevisiae*, size control primarily takes place in the G1 phase of new-born daughter cells before they are committed to cell division^7–11^. Cells that are born smaller delay entry into S phase to have more time to grow larger and compensate for their initial size^12^. In contrast, larger mother cells have homogenously short lengths of G1 phase and show no evidence of cell size control and get larger with each division cycle^7,8,13^.

The molecular network controlling the budding yeast G1/S transition has been linked to cell size regulation. Mostly, this has been attributed to the regulation of activity of the central cell cycle transcription factor SBF, a heterodimer composed of Swi4 and Swi6^14,15^. In G1, SBF is bound to and repressed by Whi5 and activated by the G1 cyclin Cln3 in complex with the cyclin-dependent kinase Cdk1 (Fig. 2A)^16–21^. In addition to Cln3 and Whi5, the gene *BCK2* has also been identified as a driver of the G1/S transition although the specific mechanism through which it works is unknown^22–24^. Once it is sufficiently active, SBF triggers a sharp positive feedback loop that commits cells to division at a point known as the *Start* of the cell cycle^10^. This feedback loop operates through the SBF-dependent transcriptional activation of the two downstream G1 cyclin genes *CLN1* and *CLN2*, which encode the Cln1 and Cln2 cyclin proteins that also form a complex with Cdk1^25^. Cln1,2-Cdk1 complexes then complete the inactivation of Whi5 to fully activate SBF and irreversibly commit the cell to division^25,26^.

While the positive feedback loop driving commitment to cell division is now well accepted, there are multiple models for how budding yeast trigger this transition. In different growth conditions, Whi5 and Cln3 concentrations likely reflect cell size and cell growth rate respectively. The concentration of Whi5 is nearly inversely proportional to cell size so that its ability to inhibit SBF decreases as cells grow larger^18,27,28^. While Cln3 concentration is nearly constant in G1 during a particular growth condition, its concentration is higher when cells consume glucose and grow quickly compared to when they consume ethanol and grow slowly^29^. While unraveling how the G1/S transition is regulated across growth conditions is an important problem, here we restrict ourselves to consider how cell size triggers an increase in the rate of G1/S progression in a given growth condition.

For the case of cell size control in a given growth condition, there remains substantial controversy in the field for how the G1/S transition is triggered in larger cells^18,30–35^. The models can divided into three groups: activator titration models^30–33,36^, inhibitor dilution models^18,37–41^, and combined models containing both mechanisms^34,35^. Activator models propose that the increase amount of activator, like Cln3 or SBF, titrate the fixed DNA amount in G1 phase. The genome is thus used as a ruler to measure the number of activator molecules, like Cln3, which would be in proportion to cell size. The inhibitor dilution model proposes that the dilution of Whi5 serves as a cell size sensor to promote the G1/S transition in larger cells^42^.

While the size control models above are consistent with the G1 cellular concentration dynamics that have been observed in budding yeast, they make different predictions for how regulatory proteins bind to the genome to control the G1/S transition. In this study, we assess cell size control models by measuring chromatin-binding kinetics of regulatory proteins in larger and smaller cells using single-molecule microscopy. We show that in daughter cells, which experience the most cell size control^7,8^, the chromatin-bound fraction of the cell cycle inhibitor Whi5 decreases as cells grow larger. This is consistent with the inhibitor dilution class of models. In contrast to Whi5, we find that the chromatin-bound fraction of the cell cycle-activating transcription factor SBF increases with cell size as cells grow during G1. Measurement of the copy numbers of these proteins in the nucleus results in an estimated ∼20 copies of SBF bound to chromatin in the smallest cells and a plateau of ∼100 chromatin-bound copies in larger cells. The transition point, where Whi5 abundance on chromatin is overtaken by that of SBF, correlates with initiation of transcription of the G1 cyclins *CLN1* and *CLN2*, as measured by single-molecule Fluorescence *In Situ* Hybridization (smFISH). This transcriptional activation of G1 cyclins marks the *Start* commitment point to the cell cycle^10,25^. Finally, we characterize protein dwell times on chromatin and show that these changes mainly result from differences in the association rate of Whi5 and SBF, as their average dwell times are ∼10 seconds across a range of cell sizes. Taken together, our data suggest a model where SBF association to chromatin, and ultimate activation of transcription, is controlled the genome-titration of SBF and the dilution of Whi5 that accompanies cell growth in G1 phase. Thus, our work marks a significant step forward towards a more mechanistic understanding of how the G1/S transition is triggered.

## RESULTS

### Single-molecule analysis of chromatin binding kinetics

To test molecular models of the budding yeast G1/S transition, we sought to measure the binding of SBF and Whi5 to chromatin in single cells. This would allow us to determine how the amount of specific chromatin bound proteins changes with cell size as cells progress through G1 phase. To do this, we made a fusion of the gene of interest at the endogenous locus with the coding sequence for HaloTag. The resulting chimeric protein was then labelled *in vivo* with cell permeable JaneliaFluor dyes (JF552 or PA-JF549)^43,44^. We used Highly Inclined and Laminated Optical sheet (HILO) microscopy^45^ to increase the signal to noise ratio in our imaging (Fig. 1A), camera integration of 10 ms to detect both diffusive and chromatin-bound molecules, and intervals of 50 ms to sample binding events lasting at least few hundreds of milliseconds. We chose these imaging parameters based on previous single-molecule imaging of yeast and mammalian transcription factors^46–50^ (Fig. 1B). Characterization of the intensity of spots and their diffusion coefficient corroborated that they are single fluorophores and that their movement is consistent with chromatin binding, resulting in a slowing down of diffusion (Fig. S1A-D). By recording the position of spots over time, we obtained maps of individual molecules (tracks), each associated with a value of their radius of gyration, as a measure of their displacement. The distribution of radius of gyrations in a population of tracks, obtained from multiple cells, were fitted to two peaks in a Gaussian Mixture Model that we interpreted as chromatin-bound and diffusive, respectively (Fig. 1C & S1). Using the information in the distribution in the population, we then calculated the fraction of molecules bound to chromatin in a single cell by categorizing tracks into either bound or unbound, and assigning them to the cells from where they originated. This allows us to correlate bound fraction with the size of the individual cell.

**Figure 1.**
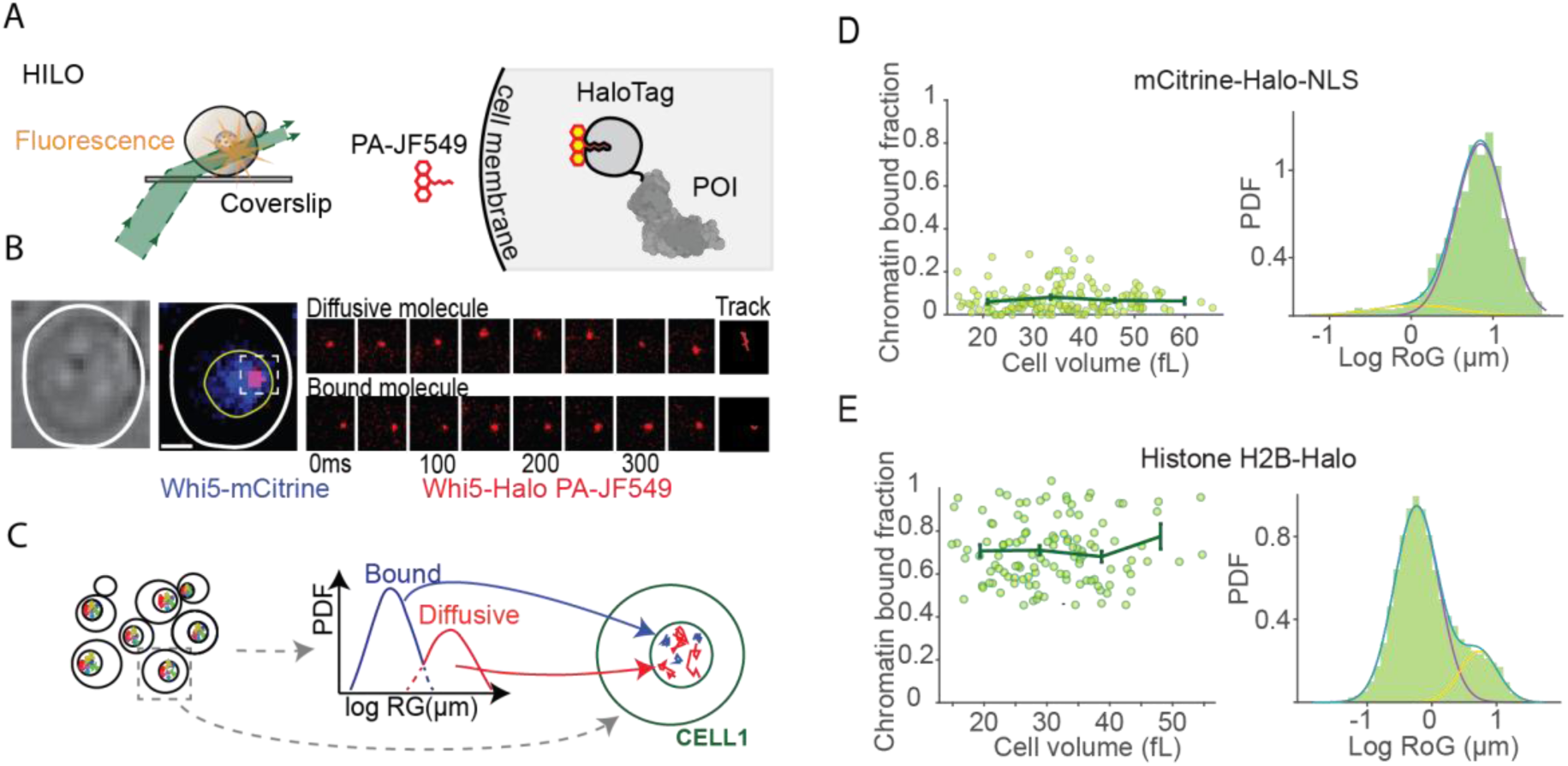
Single cell characterization of chromatin binding. (A) Proteins of interest were endogenously tagged with HaloTag and labelled with a cell permeable fluorophore coupled with a HaloTag ligand. HILO microscopy was used to increase the signal-to-noise ratio needed for single-molecule tracking. (B) Representative images of single-molecule tracking experiments. Brightfield image (left) and Whi5-mCitrine (middle) image were used to identify cells in the *pre-Start* G1 and to segment nuclei. Multiple timepoiπts of a freely diffusive molecule (top) or a chromatin-bound molecule (bottom) are shown along with the resulting ‘track’ that joins all localizations (right). (C) Procedure to characterize single cells. Tracking of molecules is done in a population of cells, resulting in a distribution of dynamic behaviours characterized by their radius of gyration. Two behaviours, chromatin-bound and diffusive, are identified in this distribution, and individual tracks, belonging to specific cells, are assigned to either of these two groups. Consequently, a fraction of chromatin-bound molecules is assigned to each cell. (D) Characterization of diffusive control mCitrine-Halo-NLS. The plot in the left compares bound-fraction against cell size. Each point in the plot represents a single cell. The plot in the right shows the log transformed radius of gyration for the cell population. (E) Single cell analysis (left) and distribution of the cell population (right) for the chromatin-bound control, histone H2B-HaloTag.

As positive and negative controls for our method of estimating chromatin bound fractions, we used a strain expressing the nuclear localized fluorescent protein mCitrine-Halo-NLS and a histone H2B-Halo fusion protein. mCitrine-Halo-NLS and H2B-Halo are expected to be largely freely diffusing or chromatin bound, respectively. As expected, molecules of mCitrine-Halo-NLS were mostly categorized as diffusive, with an average of 7.1% ± 1.1% categorized as chromatin-bound (Fig. 1D). In contrast, molecules of H2B-Halo were mostly chromatin-bound, with an average of 71.3% ± 2.5% chromatin-bound molecules per cell (Fig. 1E), similar to previous estimates of the bound fraction of histones in yeast^51^. As expected, neither of these two proteins showed any relationship between cell size and chromatin binding (mCitrine-Halo-NLS: p value = 0.984; H2B-Halo: p value = 0.689).

### Cell size increases the fraction of chromatin-associated SBF and decreases the amount of chromatin-associated Whi5 in daughter cells

To test how cell size regulates the G1/S transition, we sought to measure how the chromatin association of the key regulators Whi5 and SBF changes with increasing cell size. This is important because the activation of the Cln1,2-dependent positive feedback loop is regulated at SBF-bound promoter regions. Therefore, determining the amount of Whi5 and SBF on chromatin should give insight into their amounts at the key *CLN1* and *CLN2* promoter regions. We are particularly interested in pre-*Start* cells and how cell size drives cells through the *Start* transition. The complete dissociation of Whi5 from chromatin occurs after *Start,* when one can see its complete exit from the nucleus and the complete transcriptional activation of *CLN2*^10,25,52^.

To determine how Swi4 and Whi5 chromatin association changes with cell size, we used a strain containing either genomically integrated *WHI5-mCitrine-Halo* or *WHI5-mCitrine* and *SWI4-Halo* alleles. Both of these fusion proteins are functional since we do not observe any changes to the cell size distribution of the population, which would be expected if the fusions disrupt these proteins functions^17,18,53,54^ (Fig. S2A). As Swi6 also is also part of the transcription factor MBF^55^, we also generated a strain carrying *WHI5-mCitrine* and *SWI6-Halo* alleles in an *mbp1Δ* background. We examined this background because Mbp1 forms the MBF transcription factor together with Swi6 and we only wanted to examine SBF. As expected, this strain displayed a slight cell size increase consistent with previous results^56^(Fig. S2B). In brightfield images, we distinguish mother-daughter pairs based on morphology: the larger, more ovoid cell is classified as the mother, and the smaller, rounder cell is classified as the daughter. We identified pre-*Start* cells by the presence of nuclear-localized Whi5-mCitrine (Fig. 2B). Applying the approach described above, we quantified the proportion of chromatin-bound molecules for each cell and its size.

**Figure 2.**
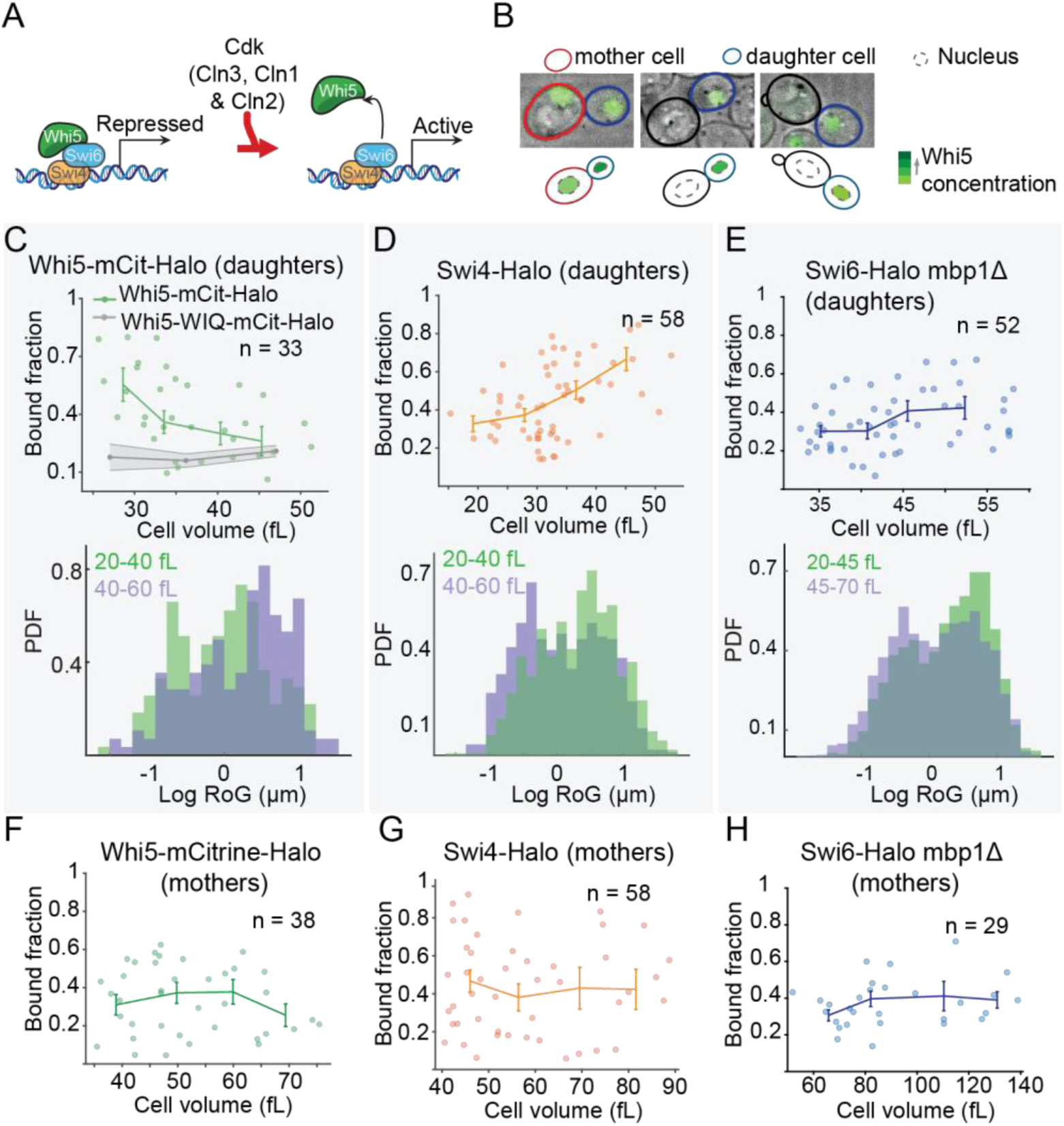
Bound fraction of SBF and Whi5 changes with size in daughter cells. (A) Diagram representing the regulation of SBF-dependent transcription during G1. (B) Representative images of the categorization done for the selection of daughter cells. Examples of overlays containing brightfield and Whi5-mCitrine images, showing pre-Sfarf daughter cells coupled to mother cells at different stages of G1 or S phases. (C) Results for the single cell analysis of chromatin binding for Whi5-mCitriπe-HaloTag (top) and for the distributions of log transformed radius of gyration for tracks obtained from small (20-40 fL) or large (40-60 fL) daughter cells (bottom). The gray line in the single cell analysis represents the trend obtained for the Whi5-WIQ mutant. (D) Results of the single cell analysis of chromatin-binding for Swi4-Halo (top) and for the distribution of radius of gyration values in small and large cells (bottom). (E) Results of the single cell analysis of chromatin-binding for Swi6-Halo mbplΔ (top) and for the distribution of radius of gyration values in small and large cells (bottom). (F-H) Results of the single cell analysis of chromatin binding for Whi5-mCitrine-Halo, Swi4-Halo or Swi6-Halo mbplΔ, respectively.

Our analysis revealed that the fraction of Whi5-mCitrine-Halo that is chromatin-bound in daughter cells was lower in larger cells (Fig. 2C; p value <0.033). For example, in cells less than 30fL in volume, 48.9% ± 9.7 % (standard error of the mean) Whi5-mCitrine-Halo is chromatin-bound, while this fraction decreased to 29.1% ± 4.4 % in cells larger than 40fL. We also examined the chromatin-bound fraction of Whi5-WIQ-mCitrine-Halo, which is defective in binding to SBF and therefore to SBF-regulated promoters^57^. The fraction of less mobile, likely chromatin-bound Whi5-WIQ is 17.4% ± 1.9 % in large and small (Fig. 2C & S3C). These results demonstrate that the fraction of Whi5 bound to chromatin decreases with cell size in daughter cells until reaching near baseline levels at ∼40fL. This baseline level was determined by analysis of the SBF non-binding mutant Whi5-WIQ-Halo. It also shows that, once Whi5 unbinds from chromatin, it can remain in the nucleus for some period of time without being immediately exported since if nuclear export immediately followed unbinding, no drop in the bound fraction would be observed and highly mobile nuclear Whi5 would be rarer.

In contrast to Whi5, both subunits of SBF, Swi4-Halo and Swi6-Halo, showed an increase in their chromatin-bound fraction as cell size increases. Small cells less than 30 fL in volume showed a bound fraction of 30.5 % ± 3.0 % of Swi4-Halo, while cells greater than 45 fL in volume showed a chromatin-bound fraction of 57.7% ± 7.4% (Fig. 2D; p value < 0.0006). For Swi6-Halo, the bound fraction increased from 21.2 % ± 3.5 % in cells <35 fL to 33.2 % ± 4.1 % in cells >50 fL (Fig. 2E; p value <0.028). Similar conclusions can be reached by analysing data from a population of cells, instead of single cells, after categorizing them into two groups (small and big cells) based on their volume (Fig. 2C-E & S3A).

It is important to note that these clear trends in chromatin binding of Whi5 and SBF are unique to daughter cells. Analysis of the chromatin-bound fraction per cell size in mother cells show no correlation, and greater scatter across cells sizes (Fig. 2F-H). This is consistent with these cells being at different stages of SBF activation in G1 irrespective of their size (Fig. S4).

### The number of chromatin-bound SBF molecules increases with cell size from tens to hundreds

To relate the chromatin-bound fraction changes we observed to absolute numbers of molecules, we measured the copy number of Whi5 and SBF molecules per nucleus in our conditions. To do this, we measured the total intensity of cells carrying mNeonGreen fusions by integrating the intensity obtained from a Z-stack obtained using a spinning disc microscope. We corrected for cell autofluorescence by collecting data with two emission filters (one for green and one for red light) but using a single excitation wavelength^58^. We then divided the integrated intensity in the cell (or the nucleus) by our estimate of the fluorescence emitted by a single-molecule. This estimate was obtained by measuring the fluorescence intensity emitted by a nuclear pore complex that has a known number of mNeonGreen labeled subunits^59^ (see methods; Fig. 3A). Our results show that Swi4 copy number scales proportionally with cell size, starting with an average of less than 50 molecules per cell in small <20FL cells and increasing to more than 125 molecules in larger >50fL cells (Fig. 3B). The abundance of Whi5 and Swi6 was higher (∼300 copies per cell), but the number of these molecules per cell increased less with cell size (Fig. 3B, Table S1). These estimates are similar to previous copy number estimates^60^, and consistent with the size-dependent concentration measurements previously reported, namely, that Whi5 is diluted in larger cells^42^.

**Figure 3.**
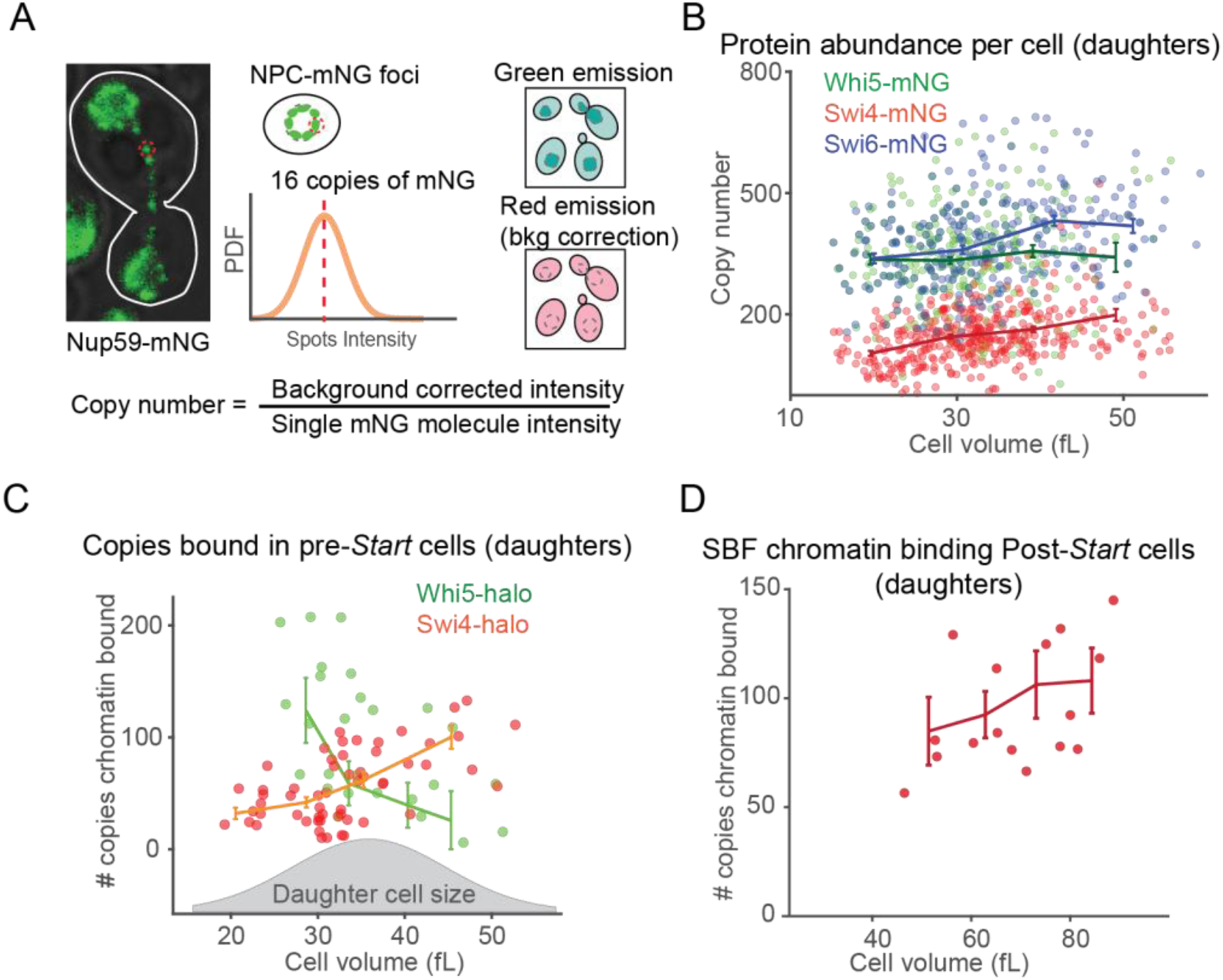
Number of chromatin-bound SBF increases with size in daughter cells. (A) Procedure used for the estimation of protein copy numbers per cell. Individual foci produced by the NPC subunit Nup59-mNeon-Green were used to characterize the intensity produced by a single mNeonGreen molecule. This value was used to divide the total cell intensity of mNeonGreen fusions, after subtracting background autofluorescence, to obtain an estimate of the copy number. (B) Distribution of copy numbers for Swi4, Swi6 and Whī5 fusions to mNeonGreen. Each dot represent a single cell. Line represents the average copy number at four different cell sizes. (C) Single-cell analysis showing the number of copies of Swi4 or Whi5 bound to chromatin with respect to the cell size for pre-Síartdaughter cells. A distribution of cell sizes is shown at the bottom of the plot. (D) Single cell analysis of the number of Swī4-HaloTag bound to chromatin in post-Sŕart daughter cells.

After measuring how protein abundance changes with cell size, we can now convert our chromatin-bound fractions to estimates of the total amount of chromatin-bound proteins. This shows that the number of chromatin-bound SBF (Swi4) molecules was 39.9 ± 3.9 per cell in small daughter cells (cell size < 30 fL), increasing to 99.8 ± 12.4 molecules per cell in large daughter cells (cell size > 45 fL). Similarly, Swi6 binding increases from 80.0 ± 13.3 molecules per cell (cell size < 35 fL) to 130.0 ± 15.9 molecules per cell (cell size > 50 fL). By subtracting the Whi5-WIQ chromatin-bound fraction, the trend is inverted for Whi5, where cells with a volume < 30 fl have 102.1 ± 32.8 chromatin-bound molecules, which drops to nearly zero in larger cells. (Fig. 3C). In comparison, post-*Start* cells have a flatter trend for chromatin-bound Swi4 starting at about the maximal occupancy observed for pre-*Start* daughter cells, suggesting a plateau in the binding at just over 100 molecules (Fig. 3D).

### Changes in chromatin binding of Whi5 and Swi4 correlate with SBF-dependent transcription

The results above show changes in the chromatin-bound number of SBF and Whi5 occurring at sizes similar to those previously associated with *CLN2* transcription and the *Start* transition^25,61^. To confirm that this is indeed the case here, we sought to measure the expression of the G1 cyclins *CLN1* and *CLN2* in single cells using single-molecule Fluorescence *In-Situ* Hybridization (smFISH) to detect mRNA transcripts of these genes in individual cells. To do this, we first generated cells carrying an MS2 24x array inserted at the 3’ end of *CLN1* and *CLN2*, fixed and permeabilize the cells, and used fluorescent probe carrying Quasar 570 fluorophore targeting the MS2 array on mRNA molecules. We then took images in the green and red fluorescent channels to detect Whi5-mNeonGreen and the MS2 array, respectively, and in brightfield to measure the cell size. To test for cells actively expressing *CLN1* and *CLN2*, we counted the number of cells that contained clear spots in the nucleus. Spots outside of the nucleus likely represent earlier transcription events (Fig. 4A). Our results showed few events in small cells, only 30.89% of 30fL cells having nuclear spots for *CLN1* (N = 123, SE = 4.2%), and an increase in the frequency of transcription events in cells larger than 35fL, with 78.54% of 40fL cells having nuclear spots for *CLN1* (N = 247, SE = 2.6%) (Fig. 4B & S5). These results corroborate our assumptions that the changes in chromatin binding correlate with changes in the expression of SBF-regulated genes in the conditions studied.

**Figure 4.**
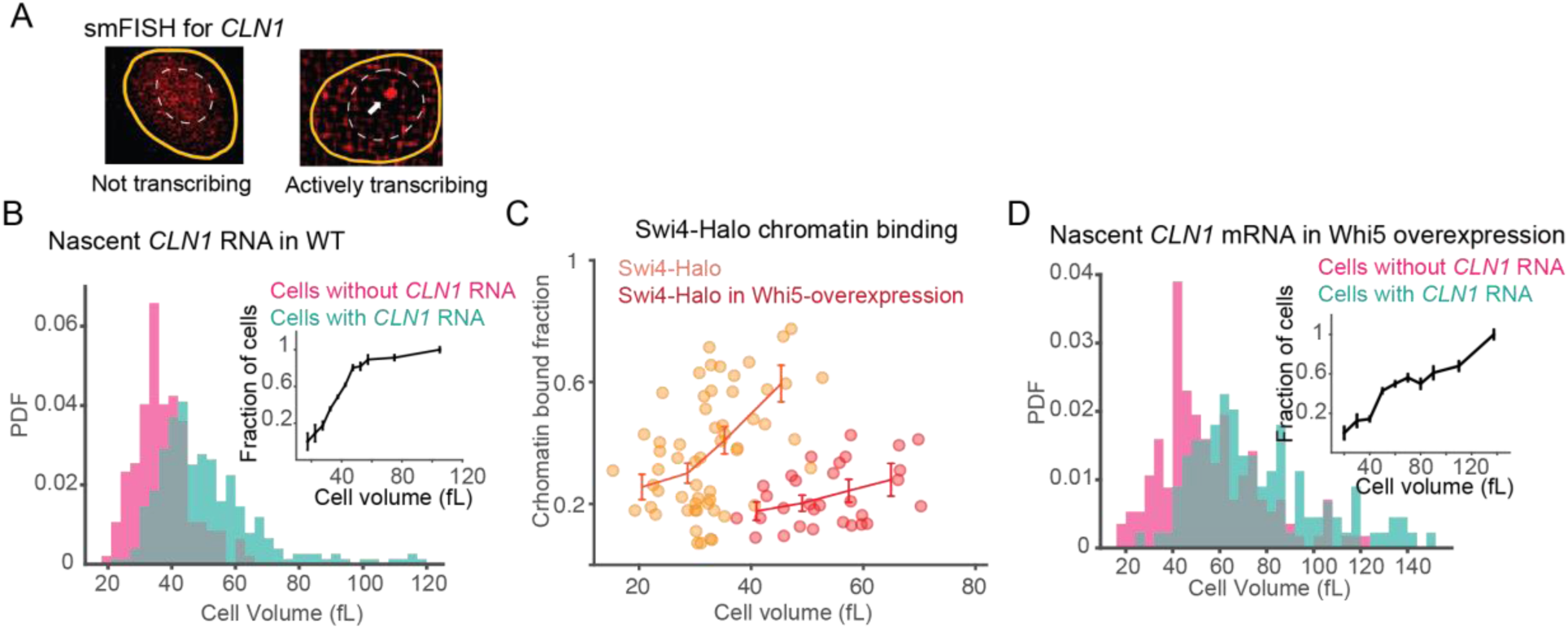
SBF-dependent gene activation matches with changes in its chromatin binding. (A) Examples of smFISH for the SBF-dependent gene *CLN1.* Cell (yellow) and nucleus outlines (white dashed) are shown. Arrow shows an example of recently transcribed mRNA molecule used for the analysis. (B) Frequency distribution of cells with (pink) or without (blue) nuclear foci representing recently transcribed *CLN1* events contrasted against cell size. Inset shows the same data but as fraction of cells with nuclear *CLN1* foci in relation with cell size. (C) Single cell analysis showing the fraction of chromatin-bound Swi4-HaloTag against cell size for wild type (orange; data identical to Fig. 3C) and for a strain overexpressíng Whi5 (red). (D) Frequency distribution of cells with (pink) or without (blue) nuclear foci of *CLN1* against cell size in cells overexpressing Whi5. Inset shows the same data plotted as frequency of cells with nuclear *CLN1* foci against cell size.

### SBF binding to chromatin is inhibited by Whi5

The observation that SBF association with chromatin increased with cell size posed a conundrum because the concentration of both its subunits, Swi4 and Swi6, did not increase with cell size (Fig. S6B). One possible explanation for the increased association of SBF with chromatin in larger cells is that Whi5 prevents SBF from associating with chromatin. Consequently, increasing chromatin-binding of SBF could reflect the lower concentration of Whi5 in larger cells. To test this hypothesis, we overexpressed Whi5 by adding a second copy of the *WHI5* gene under the control of the strong *ACT1* promoter^38^. In these cells, the bound fraction of SBF (median = 0.19) is significantly reduced compared to wild type cells (median = 0.36, p value= 0.0006). The size-dependent increase is also slightly flattened compared to the wild type cells (p value= 0.09) (Fig. 4C). smFISH data confirmed that in cells overexpressing Whi5, the expression of *CLN1* generally occurs at cell sizes larger than 40 fL (compared to 30 fL in WT), consistent with the changes in binding of SBF we observe (Fig. 4D). Taken together, these data indicate that SBF binding is regulated by Whi5.

### Dynamic association of SBF to chromatin is hindered by its interaction with Whi5

Having found that SBF is less chromatin-associated in Whi5 over-expressing cells, we next sought to determine why this is. In principle, it could be from reducing the rate free SBF binds chromatin or the time it stays bound there. We therefore sought to characterize the dwell time of SBF and Whi5 using single-molecule microscopy. Previous work using chromatin immunoprecipitation has demonstrated binding of SBF and Whi5 at promoters during G1^16,17^, but the dwell time associated with this chromatin-binding has not been determined. To measure dwell times on chromatin, we used an approach similar to that we previously applied to characterize the subunit dynamics at the replisome^62^. Briefly, as in the bound-fraction experiments described above, we used HaloTag derivatives labelled with PA-JF459 dye and HILO microscopy. The change to the protocol is that we now used camera integration times of 500 ms rather than 10 ms. This longer acquisition time resulted in motion blurring of fast-moving molecules but not chromatin bound molecules (Fig. 5A; see methods). We identified chromatin-bound molecules using a machine-learning approach^63^, and inferred the time a molecule was associated with the chromatin from the lifetime of the spots representing bound molecules after correcting for photobleaching (Fig. 5B & S7). Histone H2B-Halo was used as our bleaching control, as we expect that its focus disappearance largely represents bleaching rather than unbinding from the chromatin.

**Figure 5.**
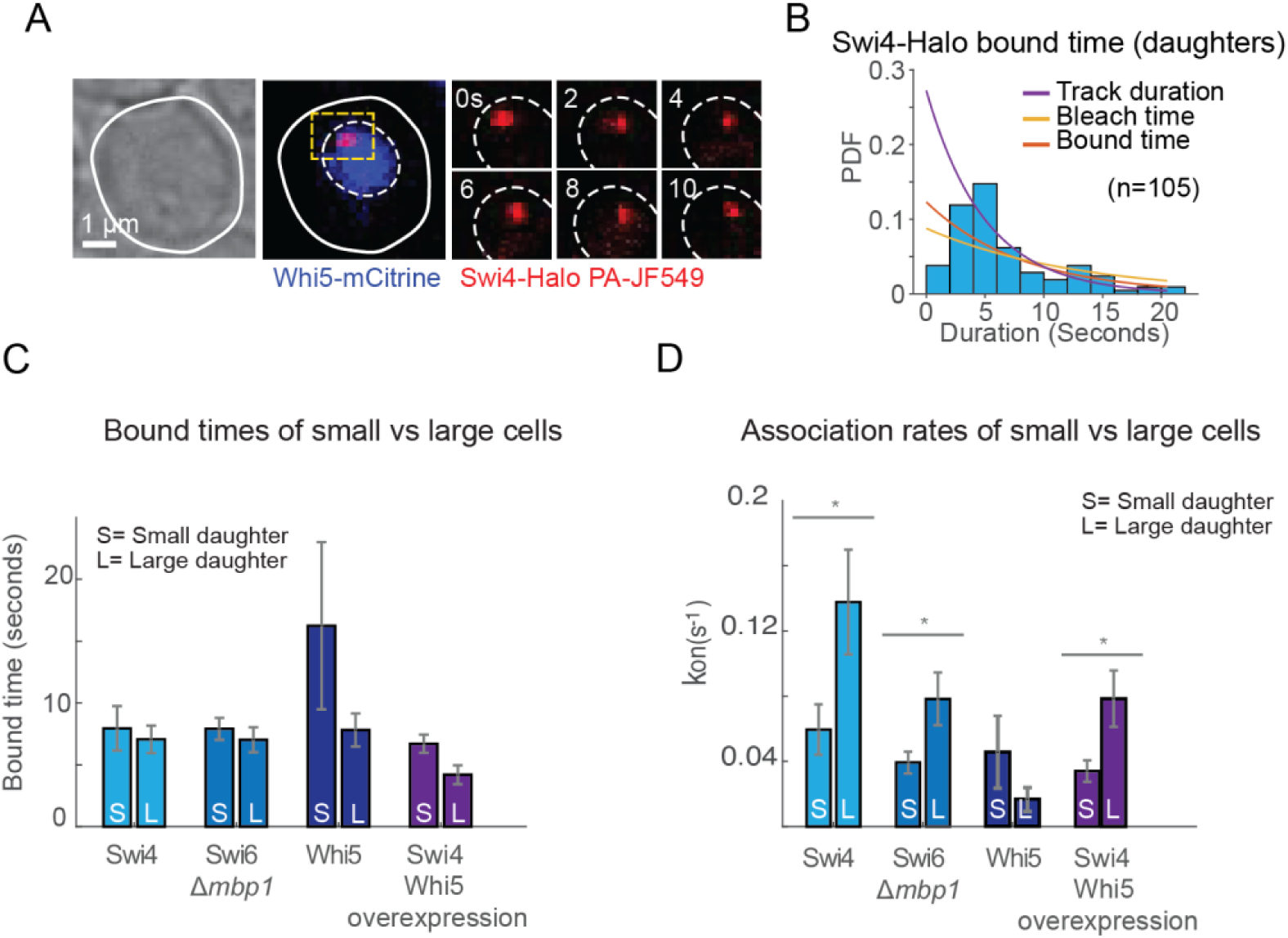
SBF binding to chromatin is transient and is inhibited by Whi5. (A) Representative images of the imaging done to estimate dwell times. A brightfield images (left) is used to determine the cell outline. Whi5-mCitrine (middle) marks pre-Start cells and is also used to segment nuclei. Images of a chromatin-bound Swi4-HaloTag coupled to PA-JF549 are shown at different time points. (B) Example of a distribution of lifetimes for 105 Swi4-HaloTag foci (blue bars). The distribu­tion is fit to a single exponential decay (purple), which after correcting it by the bleaching control (yellow), results in a corrected distribution of bound times (orange). (C) Estimated dwell time values for SBF and Whi5 in daughter cells categorized as small or large. Error bars represent standard error of the mean. (D) Estimated association rate values for SBF and Whi5 in daughter cells categorized as small or large. Error bars represent standard error of the mean.

Results from daughter cells show that Swi4 has an average dwell time of 8.1 seconds (C.I. 5.9-10.6 s), Swi6 has a dwell time of 12.2 seconds (C.I. 9.4-15.5 s), and Whi5 has a dwell time of 9.6 seconds (C.I. 6.9-12.7 s) (Fig. S7C). It is interesting that dwell time estimates are similar for SBF and Whi5 because this suggests most chromatin binding and unbinding of Whi5 occurs when it is associated with SBF. The duration of chromatin association of SBF does not differ significantly between daughters and mother cells (Fig. S6C) or between small and large daughter cells (Fig. 5C). Moreover, the average dwell time of SBF is similar in Whi5 overexpressing cells (Fig. 5C & S7), suggesting that Whi5 does not affect the dissociation of SBF from its binding sites on the genome. In contrast, estimates of the association rates, calculated using the measured bound fraction and dwell times, show a significant difference in small and large cells (Fig. 5D). These results show that regulation of transcription of SBF occurs by transient binding to promoters. Furthermore, since Whi5 affects the average occupancy of SBF on the genome, this result implies that forming a complex with Whi5 reduces the rate SBF binds its target sites on the promoter, and not the time it spends on the promoter after binding.

## DISCUSSION

Our live cell study of SBF and Whi5 chromatin association gives mechanistic insight into how cell size triggers progression through the *Start* transition in budding yeast. As daughter cells grow larger, less Whi5 is associated with the chromatin. This is consistent with the inhibitor dilution model where the decreasing concentration of Whi5 in larger cells promotes cell cycle progression^42^. In addition to linking Whi5 dilution to changes in chromatin occupancy, we observe that SBF binding to chromatin increases with cell size, despite its approximately constant cellular concentration. This is consistent with SBF titration contributing to the size-dependence on *Start*. Moreover, it is likely these two mechanisms are intertwined as Whi5 inhibits SBF association with the chromatin (Fig. 6).

**Figure 6.**
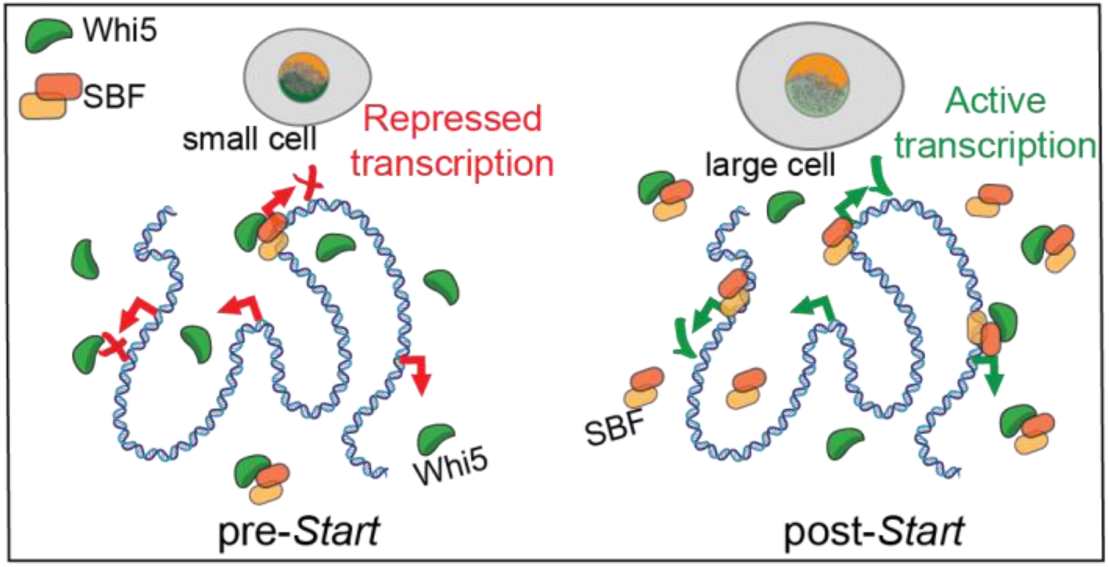
Model for SBF regulation in daughter cells. Small newly bom daughter cells have a few-fold excess of Whi5 relative to SBF (left). Whi5 limits SBF binding to chromatin and represses SBF-dependent transcription. As cells grow, Whi5 is diluted and the number of SBF copies increases (right). This shift in relative abundance increases the fraction of SBF not bound to Whi5, promoting SBF binding to target promoters and ultimately activating SBF-dependent transcription.

Our data on the absolute numbers of molecules bound to chromatin has additional implications for the regulation of cell cycle-dependent gene expression. We estimate that the number of chromatin-bound Swi4 molecules starts at about 40 in small G1 cells and rises to about 100 as cells arrive at *Start*. Given that there are approximately 200 G1/S transition-related promoters, and that some of these promoters contain multiple SCB sites^64–66^, not all SBF-regulated promoters can be bound to SBF even when they are all active. This conundrum is resolved once we consider the rate with which SBF interacts with chromatin. Our data show that both Whi5 and SBF have a similar dissociation rate on chromatin, with average dwell times of ∼10 seconds. This is within an order of magnitude to the reported dwell times of other yeast transcription factors such as Ace1p, with a ∼3 second average^67^, and Gal4, with a reported average of 25-75 seconds^68^. These estimates are also within the range of transcription factor dwell times reported for mammalian cells, which span from a few to hundreds of seconds^69^. The fast turnover of SBF on chromatin means that a smaller number of transcription factors can regulate a larger number of binding sites on the genome. This is because SBF will be bound to each of its sites a fraction of the time depending on its specific binding constants so that the same molecule can regulate several sites in any given minute.

Our data show that changes in SBF and Whi5 binding to chromatin are largely explained by changes in their association rates rather than their dissociation rates. This suggests that Whi5 primarily reduces the probability of SBF associating with chromatin, rather than actively promoting its dissociation. Together with the matching dwell times of SBF and Whi5, and prior evidence for their direct interaction in solution^16,17^, these results support a model in which SBF diffuses and binds to chromatin as a complex with Whi5. Reduced chromatin binding of SBF, but not its complete abrogation, suggests that either a minimal number of copies of SBF binding is required for promoter activation, or that the presence of Whi5 prevents promoter-activation. Changes in the binding of Whi5 may at least partly reflect changes in its phosphorylation state^70^, as phosphorylated Whi5 is unable to bind chromatin through SBF. Further work will be needed to determine whether Whi5 dissociation from chromatin precedes its phosphorylation or simply reports on it.

Our quantitative analysis has highlighted the need for additional biochemical studies of the Whi5-SBF interaction. We found that there is more Whi5 bound to chromatin than Swi4 in the early G1 phase. However, it was previously shown that Whi5 ChIP signal depended on its GTB region that likely interacted with Swi6’s C-terminal domain^57^. Consistent with these results, we also found that the WIQ *WHI5* mutation disrupts its chromatin association. This raises the question that if Whi5-SBF were a simple protein-protein interaction, and Whi5’s chromatin association depended on SBF, then how could more Whi5 than SBF molecules be bound to chromatin at any time? One possibility is that SBF has multiple binding sites for Whi5. Another is that Whi5 may bind to another copy of Whi5 that is bound to SBF. Additional research is required to resolve this intriguing mechanism.

Taken together, our findings based on live cell single-molecule microscopy provide a mechanistic framework for understanding how cell size regulates cell cycle progression through the G1/S transition in budding yeast. We demonstrate that SBF association with chromatin increases with cell size, while Whi5 association decreases, illustrating the intertwined dual contributions of activator titration and inhibitor dilution to cell cycle regulation. These results bridge the gap between known concentration dynamics and the biochemistry taking place at the promoters driving commitment to the *Start* of the cell cycle. Moreover, our quantitative analysis of the dynamic interplay between Whi5 and SBF on chromatin identify new questions related to their biochemical mechanism. Beyond yeast, these insights provide a conceptual framework that may inform our understanding of size control and cell cycle regulation in multicellular organisms, including humans, where similar principles governing cellular growth and division have been shown to apply.

## Acknowledgments

We thank Janelia Farms for sharing the HaloTag ligand conjugated with PA-JF549 and JF552 with us. We thank other members of the Reyes and Skotheim Labs for helpful discussions. RRL was supported by the Natural Sciences and Engineering Research Council of Canada (RGPIN-2019-05701), Canadian Institutes for Health Research (PJT-162247 and PJT-178397), the Canada Foundation for Innovation (#228994) and the Canada Research Chairs program. JMS group was supported by NIH R35 GM134858.

## Resource Availability

Requests for further information and resources should be directed to and will be fulfilled by the lead contact, Rodrigo Reyes. All unique reagents generated in this study are available from the lead contact without restriction. Scripts used are available at https://github.com/thereyeslab/General-scripts

## Supplementary Information

Document S1. Figures S1-8 and Tables S1-4

## SUPPLEMENTARY INFORMATION

### METHODS

#### Strains and constructions

Most *S. cerevisiae* budding yeast strains used in this study are in the W303 background (Table S1). Full genotypes of all strains used in this study are listed in Table 1. The plasmids used in this study are based on pUC18, which has a ColE1 origin and ampicillin resistance. The plasmid pTB16 contains the mNeonGreen gene followed by a NatMX marker, while pSJW01 includes the HaloTag gene with a HygB marker. Both the mNeonGreen and HaloTag genes contain an 8 amino acid linker sequence (GGTGACGGTGCTGGTTTAATTAAC) at the 5’ end. For insertion of an extra copy of Whi5 in the *URA3* locus, plasmid pMS192 and derivatives were cut with PacI and the digest was transformed into the yeast strain. These plasmids were maintained in E. coli DH5α, cultured in LB medium with 100 μg/ml ampicillin, and extracted using the Presto Mini Plasmid Kit from Geneaid.

All PCRs were conducted using Q5 polymerase from NEB. Each PCR reaction had a total volume of 50 μl, consisting of water, reaction buffer, 2.5 mM of each dNTP, 0.2 μM of each primer, 1 ng of plasmid DNA for insertions or 1 μl of genomic DNA for screening insertions, and 0.5 μl of polymerase.

Fluorescent fusions were generated by PCR amplification from pTB16 or pSJW01. The PCR products were transformed into haploid W303 using the following protocol. A single colony was cultured overnight at 30°C in 5 ml yeast peptone dextrose (YPD), then diluted to an OD600 of 0.1 in 10 ml of YPD. Cells were collected at an OD600 of 0.5-0.6, centrifuged at 4000 rpm for 5 minutes, and the pellet was washed twice with 25 ml of sterile deionized water and once with 1 ml of 100 mM lithium acetate. Cells were resuspended in 240 μl of 50% PEG, followed by the addition of 50 μl salmon sperm DNA (heated at 95°C for 5 minutes, then cooled on ice for at least 10 minutes), 50 μl PCR product, and 36 μl of 1M lithium acetate. The mixture was pipetted thoroughly and incubated on a rotator at 25°C for 45 minutes, then heat-shocked at 42°C for 30 minutes. The cells were pelleted using a microcentrifuge, washed with 500 μl of sterile water, resuspended in 200 μl of YPD, and plated on YPD agar. After overnight incubation at 30°C, the cell lawn was replica-plated onto selective YPD agar with 100 μg/ml NAT (for mNeonGreen) or 200 μg/ml Hygromycin B (for HaloTag). Transformants were screened for inserts via PCR.

Generation of strains carrying MS2 fusions to *CLN1* and *CLN2* to detect corresponding mRNA for smFISH experiments were generated by using the plasmid pET264 (1) to amplify 24xMS2V6 along with homologous arms for insertion at the 5’ end of the coding sequence of the *CLN1* and *CLN2* loci, followed by standard yeast transformation and selection.

#### Strain growth for confocal microscopy

Strains were stored at –80°C after construction. They were streaked onto yeast peptone dextrose (YPD) plates and incubated at 30°C for 2 days. For each microscopy experiment, a single colony was picked and grown in 5 mL SC medium with shaking at 30°C for over 8 hours. The culture was then diluted in 5 mL SC to an OD600 of 0.0047 for another 12 hours incubation. Cells were diluted again to an OD600 of 0.15 the next day, and further grown until the OD600 reached 0.3. For confocal microscopy, cells were washed once in 500 μL of SC and spread on a 1% agarose pad in SC for imaging.

#### Single-Molecule Imaging

A single yeast colony from a YPD plate was placed in 5mL synthetic complete (SC) medium and grown with shaking at 30°C for ∼7-8 hours. This culture was diluted into 5mL of fresh SC and grown overnight at 30 degrees. The overnight culture was diluted once more to an optical density at 600nm (OD600) of 0.15 the next day and grown until the OD600 reached 0.30. 1mL of this culture was spun down for 1 min at 4000 RPM, and the pellet was resuspended in 500 μL of fresh SC. Janelia Farms photoactivatable JF549 (PA-JF549) was added to the 500μL culture for a final dye concentration of 75nM, except for MS162 (Histone HTB1-Halo), where a concentration of 10nM was used to compensate for the higher copy number of histone. Cultures were placed in a thermomixer at 30°C and mixed at 550 RPM for 40 minutes. After incubation, 3 wash cycles using fresh SC were done to wash away unbound dye. After the final wash step, the pellet was resuspended in 20μL of SC, and 4μL of the culture were placed on an agarose pad consisting of SC and Optiprep (Sigma), within a Gene Frame (Thermo Scientific). The pad was made by taking a 2% agarose Optiprep mixture (0.02g in 1mL Optiprep) – that was heated to 90 degrees – and mixing 500μL with 500μL 2xSC, resulting in a 1% agarose 30% Optiprep SC mixture. Approximately 110μL of this mixture was placed within the Gene Frame, with excess being removed with a KimWipe. Prior to imaging, the sample was left for ∼15 minutes to let any unbound dye be released from cells.

Coverslips were cleaned with the following steps: 1) 2% VersaClean detergent solution overnight. 2) Washed with MilliQ water 3 times. 3) Sonicate in acetone for 30 minutes. 4) Wash with MilliQ water 3 times. 5) Washed in methanol and flame excess from coverslips using Bunsen burner. 6) Place in Plasma Etch plasma oven for 10 minutes.

Microscopy was done at 23°C on a Leica DMi8 inverted microscope with a Roper Scientific iLasV2 (capable of ring total internal reflection fluorescence (TIRF)), and an Andor iXon Ultra 897 EMCCD camera. An Andor ILE combiner was used, and the maximum power from the optical fiber was 100mW for the 405nm wavelength, and 150mW for the 488nm and 561nm wavelengths. The iLasV2 was configured to do ring highly inclined and laminated optical sheet (HILO), for selective illumination and single-molecule sensitivity. Metamorph was used to control acquisition. A Leica HCX PL APO 100x/1.47 oil immersion objective was used, with 100nm pixel size. Z stacks were done using a Plano piezo Z controller. Single-particle photoactivated localization microscopy (sptPALM) experiments were performed by using continuous activation of molecules with low power (0.1% – 10% in software) 405nm light to photoactivate ∼1 molecule/cell, with simultaneous fast-exposure (10ms) illumination with 561nm light (70% in software) to image molecules. A 10 μm Z-stack of bright field and a 561nm excitation (0.5μm step size) were acquired, to classify the G1 phase cells, estimate the cell volume, and to filter out the tracks that were not in nuclear regions.

#### Single-Molecule Tracking Analysis

Aside from tracking, all analysis was done using custom-made code in MATLAB.

Tracking was done using Trackmate (Tinevez *et al*., 2017). First, molecules were localized in each frame using a Laplacian of Gaussian (LoG) method, with an estimated diameter of 5 pixels. An intensity threshold was chosen that was slightly low, to still detect molecules that were moving out of the focal plane and were fast diffusing. After localization, tracks were formed using the Linear Assignment Problem algorithm by linking molecules in consecutive frames. The linking distance was set to 5 pixels. A gap frame of 3 was used to allow for missed localization. The gap-linking distance was set to 5 pixels more than the linking distance. Linking also had cost of 0.3 for the ‘‘Quality’’ parameter to ensure that correct molecules were linked. Tracks with less than five localizations were discarded as they were unreliable.

Cells in G1 phase were selected according to their morphology. The bright field in focus images of cells were extracted from z-stacks. Cell outlines were segmented by a custom-made MATLAB code, then modified manually by using the “Freehand” function in MATLAB. Cell volumes were estimated from the cell masks. We assumed budding yeast cells in G1 are ellipsoids. The volumes of cells were estimated by adding up the cross-section volume at each orthogonal pixel layer of the major axis of cells.

Maximum intensity projections (MIP) of z stacks in 488nm were used for segmenting the nuclear regions. Since all the proteins used in locate inside the nuclear in G1 phase, we only kept tracks that were in nuclear region. Based on the binary masks of nuclear regions, tracks that locate outside the nuclear were discarded.

Radius of gyrations (RoG) were then calculated for tracks in G1 phase cells by using the following equation:

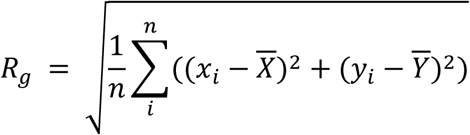

where n is the number of localizations in each track, *x*_*i*_ and *y*_*i*_ are the coordinates of the track for each step, 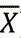 and 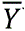 are the mean of the x, y coordinates of the track, respectively.

To classify the status of molecules, one dimensional Gaussian mixture model (GMM) was then used to fit the log-transformed RoG distribution. The initials of fitting were inferred from the log-transformed RoG distribution of H2B-Halo and mCitrine-Halo-NLS, which have most molecules bound and diffusive, respectively. To determine how many Gaussian groups exist in the RoG distribution, the distributions were fitted with different GMM models, where total gaussian groups varied from 1 to 4. The Bayesian information criterion (BIC) was calculated for each GMM fitting. The model with the lowest BIC was selected. The Gaussian group with the lowest average RoG value was classified as the group of bound molecules. The tracks inside this bound group were then marked. The chromatin bound fraction of the protein of interested in each cell were determined by dividing the number of bound tracks by the number of total tracks. Cells with total track number below 30 were discarded. Combined with the cell volume and chromatin bound fraction data of cells, we then obtained the trend of chromatin bound fraction of POIs over cell size in G1 phase.

#### smFISH sample preparation

The smFISH technique was used to investigate gene expression in yeast. Yeast strains were initially cultured in 3 mL of YPD medium at 25°C overnight in shaker. The next morning, cultures were diluted to an OD600 of 0.2-0.25 and grown to an OD600 of 0.5-0.6 in approximately 3 hours. During this time, microscope coverslips (18 mm) were coated with 0.01% poly-L-lysine, washed, and dried. Yeast cells were fixed by adding paraformaldehyde to a final concentration of 4% and incubating with rotation at room temperature for 45 minutes. The fixed cells were then centrifuged, washed twice with ice-cold 1X Buffer B, and resuspended in a spheroplasting buffer containing beta-mercaptoethanol, VRC, and lyticase. Cells were incubated at 30°C for approximately 2 minutes to achieve partial spheroplasting, checked under a microscope. Next, cells were attached to poly-L-lysine coated coverslips and incubated at 4°C for an hour.

Coverslips with attached cells were treated with 70% ethanol at –20°C overnight to facilitate probe entry. On the day of hybridization, coverslips were washed with 2x SSC and pre-hybridized with a solution of 20% formamide and 2x SSC for at least 30 minutes. smFISH probes were dried in a vacufuge and then resuspended in a hybridization buffer. The hybridization mixture was applied to the cells, and coverslips were incubated at 37°C for 3 hours. The MS2V6WT probe coupled to a Quasar 570 fluorophore was used as described previously (1).

After hybridization, coverslips were washed with pre-warmed pre-hybridization solution and 2x SSC. Finally, coverslips were dried with 90% ethanol, mounted onto microscope slides with ProLongGold, and cured overnight. The following day, coverslips were sealed with nail polish and imaged using fluorescence microscopy.

#### smFISH imaging and analysis

Samples were imaged on a Leica DMi8 inverted microscope with a Roper Scientific iLasV2, and an Andor iXon Ultra 897 EMCCD camera. A Leica HCX PL APO 100x/1.47 oil immersion objective was used, with 100nm pixel size. A series of optical sections with z-steps of 0.22 μm were collected. For GFP, an excitation line of 488 nm was used and emission at 410-540 nm. Outlines were generated using Cellpose2 (2), and single mRNA foci were labelled manually.

#### Copy number estimation

The copy number estimation was done on a Leica DMi8 inverted confocal microscope at room temperature (23°C) fitted with a Leica HCX PL APO 100x/1.47 oil immersion objective and a DISKOVERY multi-modal imaging system with a multi-point confocal 50μm pinhole spinning disk. Images were captured using dual Andor iXon Ultra 897 EMCCD cameras. Z stacks were generated with the ASI three-axis motorized stage controller and the MCL nano-view piezo stage. Image acquisition was controlled using Metamorph software. Z stacks of 15 steps (0.59 μm per step) were first captured in BrightField. Then, Z stacks of 10 steps (0.59 μm per step) were taken in the dual channel having a cut-off at 561 nm. Excitation with the 488nm laser (set to 25% in Metamorph) allowed us to simultaneously capture signal in the green and red emission channels. The exposure time for all confocal microscopy images was 200ms.

Estimation of the single-molecule intensity equivalent was done using a subunit of the nuclear pore complex as a known stoichiometry reference. Single Nup59-mNeonGreen foci were analyzed using a 2D Gaussian fit with the GaussFit On Spot plugin in ImageJ. The intensity values of individual spots were modeled with a Gaussian Mixture Model, showing two components. The mean intensity of the second component was roughly twice that of the first, aligning with the expectation that these represented single and double NPCs, respectively. Consequently, the mean intensity of the first component was interpreted as representing 16 mNeonGreen molecules.

To obtain copy number estimates, cell regions were segmented manually. G1 daughter and mother cells were selected based on the morphology of the cells. To estimate the fluorescence intensity of the protein of interested, fluorescence images were background-subtracted using ImageJ software. For each image, the background subtraction function was applied with a radius set to 20 and the smoothing option disabled. The integrate fluorescence intensity of the proteins were estimate based on the sum of each Z slice which has the protein fluorescence signal. The z range was determined by the thingSpeakRead MATLAB function to determine changes in signals using change-point detection. To remove autofluorescence background, the intensity was then normalized for the average mean value in cells that do not carry the fluorescent protein (intensity under red channel).

The copy number of the proteins per cell was estimated by using the sum intensity of the proteins divided by the intensity per mNG molecule. The concentration of the proteins was determined by using the copy number divided by the cell volume.

#### Whi5-mCitrine timelapse imaging

Whi5-mCitrine timelapse imaging was performed using an Olympus IX83. Samples were illuminated with and X-Cite 120LED lamp (Excelitas Technologies) using a YFP filter, with 500ms exposure, at an interval of 5 minutes over a total duration of 3 hours. Brightfield images were taken together with the YFP channel. Daughter cells and mother cells were separated based on the Whi5 nuclear localization and cell morphology in G1 phase in brightfield. Cell segmentation was done manually. Cell volume was estimated based on the 2D segmentation of the cell. The time Whi5 stayed in nucleus after cell division was recorded.

**Figure S1.**
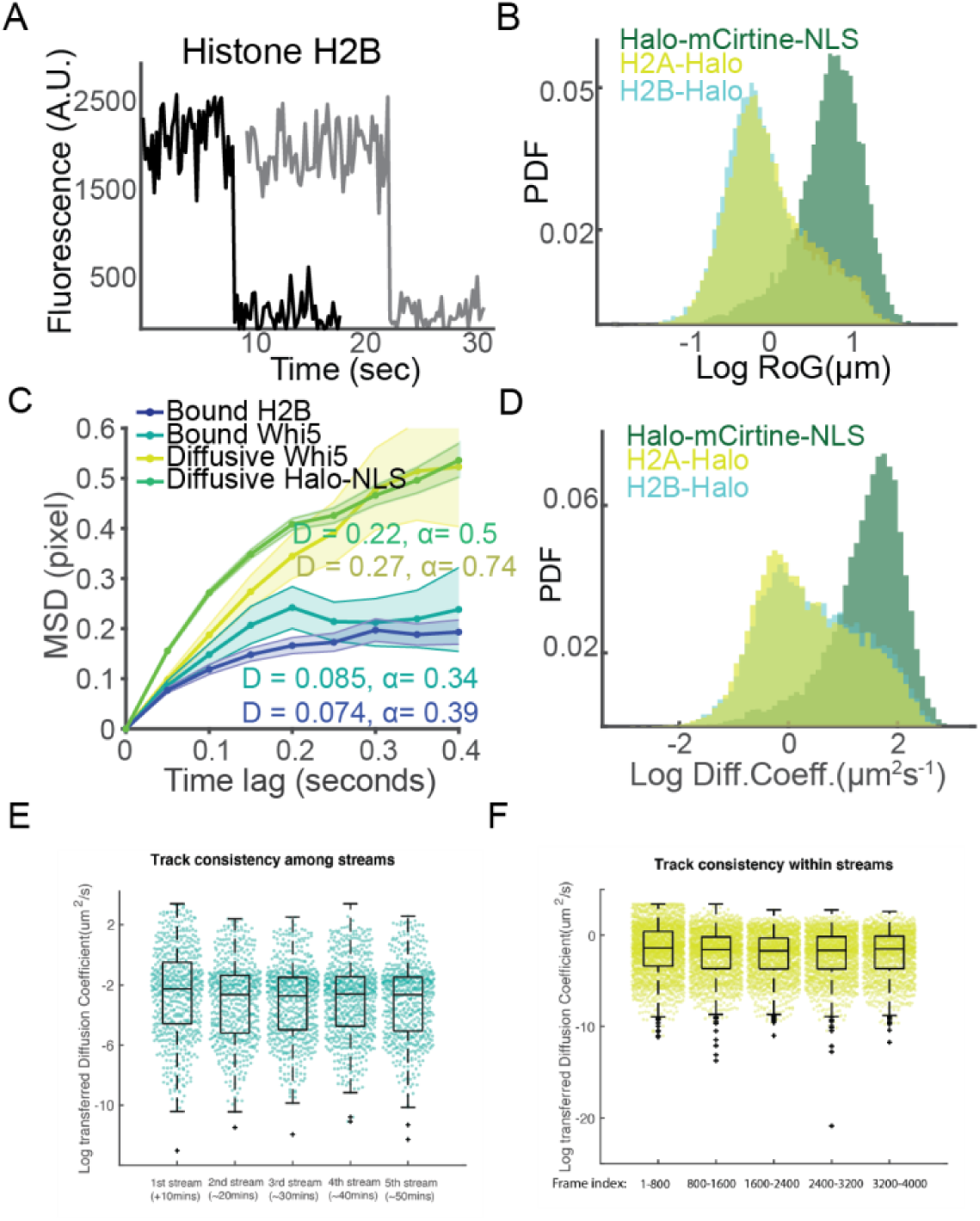
Validation and classification of chromatin-bound single-molecule tracks. (A) Representative fluorescence intensity-time traces of single H2B-Halo molecules. Both traces show a distinct single-step photobleaching event, confirming that each trace corresponds to an individual protein molecule. (B) Ensemble mean-square displacement (MSD) curves of chromatin-bound histone, chromatin-bound Whi5, diffusive Whi5, and diffusive Halo-NLS tracks over lag time. Diffusion coefficients are fitted to the MSD values. (C) Distributions of radius of gyration (RoG) and diffusion coefficients for H2B-Halo (HTB1; n = 35,978 tracks), H2A-Halo (HTA1; n = 20,034 tracks), and Halo-NLS (n = 8,057 tracks). (D) Track-to-track consistency comparison across consecutive imaging streams. (E) Iπtra-stream consistency of diffusion estimates across different time intervals within a single stream. Each boxplot represents one stream (D) or a subsegment within it (E).

**Figure S2.**
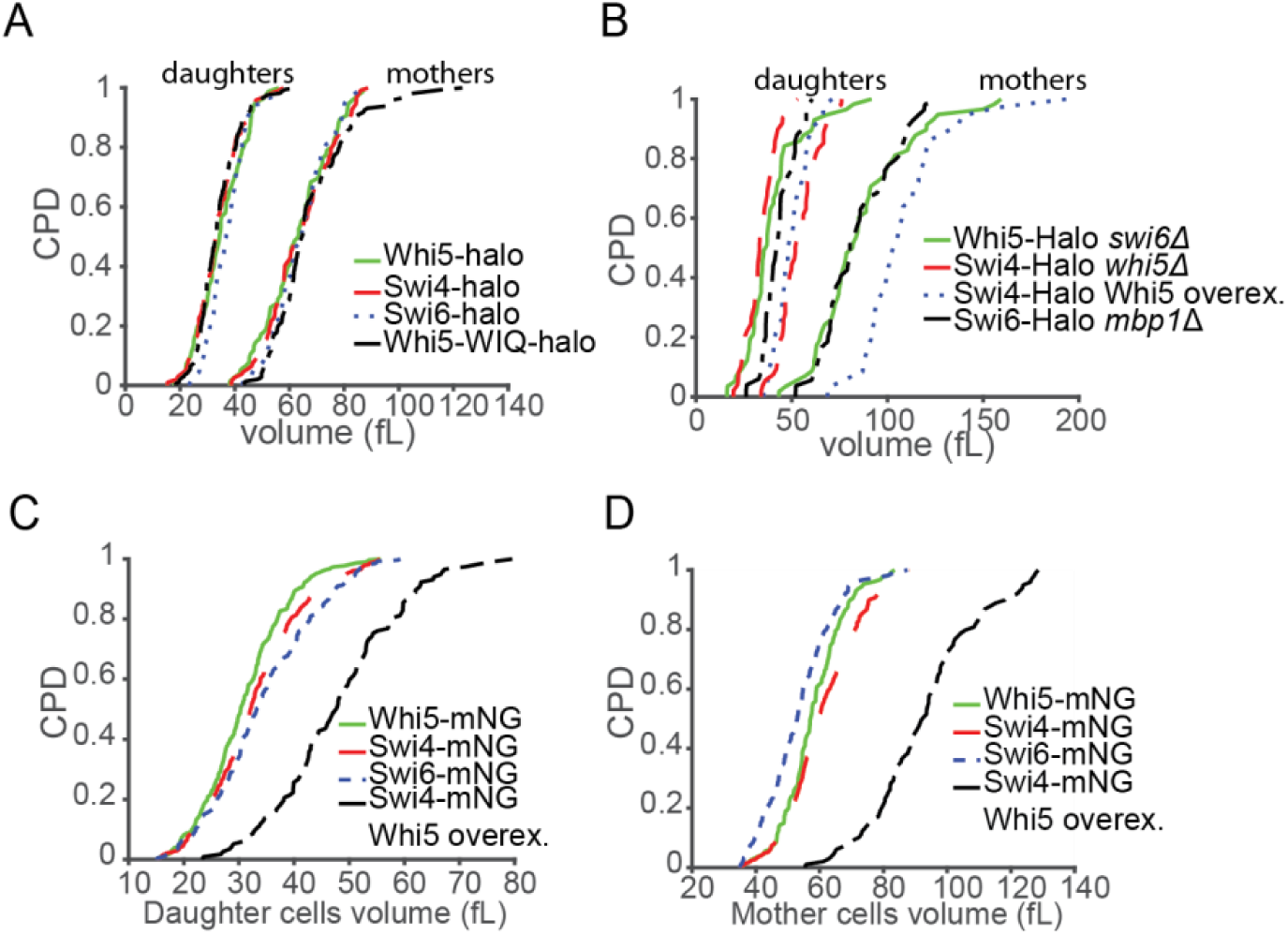
Cell-volume distributions of the yeast strains used in this study. (A) Cumulative probability distributions (CPDs) of G1 daughter (left) and mother (right) cell volumes for strains expressing Whi5-Halo (green solid), Swi4-Halo (red solid), Swi6-Halo (blue dotted), and Whi5-WIQ-Halo (black dashed). (B) CPDs for mutant and perturbed backgrounds: Whi5-Halo swi6Δ (green), Swi4-Halo whi5Δ (red dashed), Swi4-Halo with Whi5 overexpression (blue dotted), and Swi6-Halo mbplΔ (black dashed). (C) CPDs of G1 daughter-cell volumes for Whi5-mNeonGreen (mNG, green), Swi4-mNG (red), Swi6-mNG (blue), and Swi4-mNG with Whi5 overexpression (black dashed). (D) CPDs of G1 mother-cell volumes for the same mNG strains as in (C). CPD, cumulative probability distribution.

**Figure S3.**
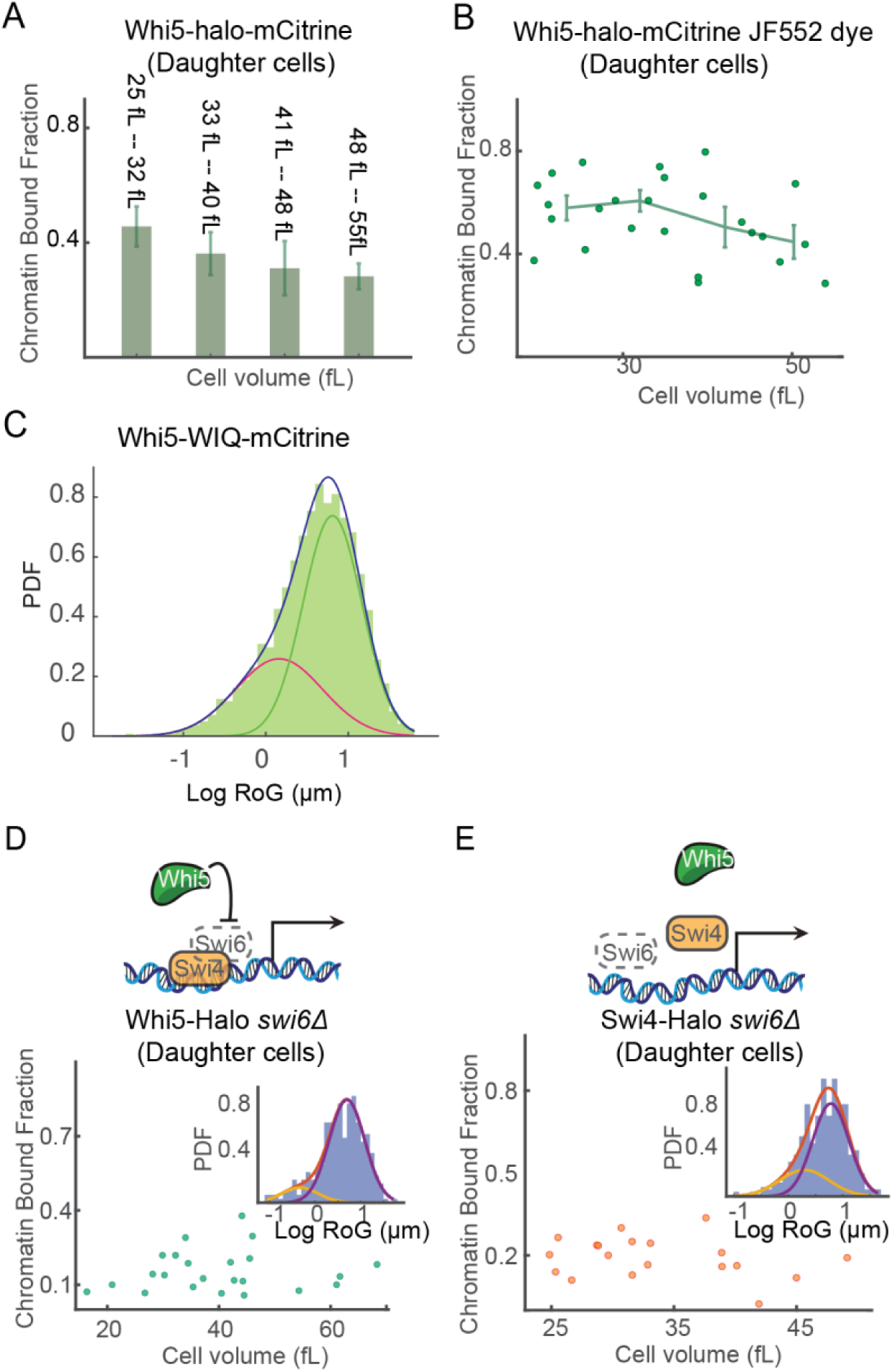
Independent dye validation and Swi6-dependence of chroma-tin-binding measurements. (A) Mean chromatin-bound fraction of Whi5-Ha-lo-mCitrine in G1 daughter cells binned by cell volume: 25-32, 33-40, 41-48, and 49-55 fL (mean ± s.e.m.). (B) Replicate measurement using the orthogonal dye JF552. Dots represent single cells; grey line indicates mean ± s.e.m. (n = 25). (C) Probability density function (PDF) of log-transformed RoG for Whi5-WIQ-mCitrine. The histogram (green) is fitted with a two-component Gaussian mixture (blue line; individual components in red and green). (D) Chromatin-bound fraction of Whi5-Halo in swi6Δ G1 daughter cells (n = 23). Inset: RoG distribution and two-component Gaussian fit. (E) Same as (D), but for Swi4-Halo in swi6Δ background (n = 18). Inset: log-RoG distribution and Gaussian fit.

**Figure S4.**
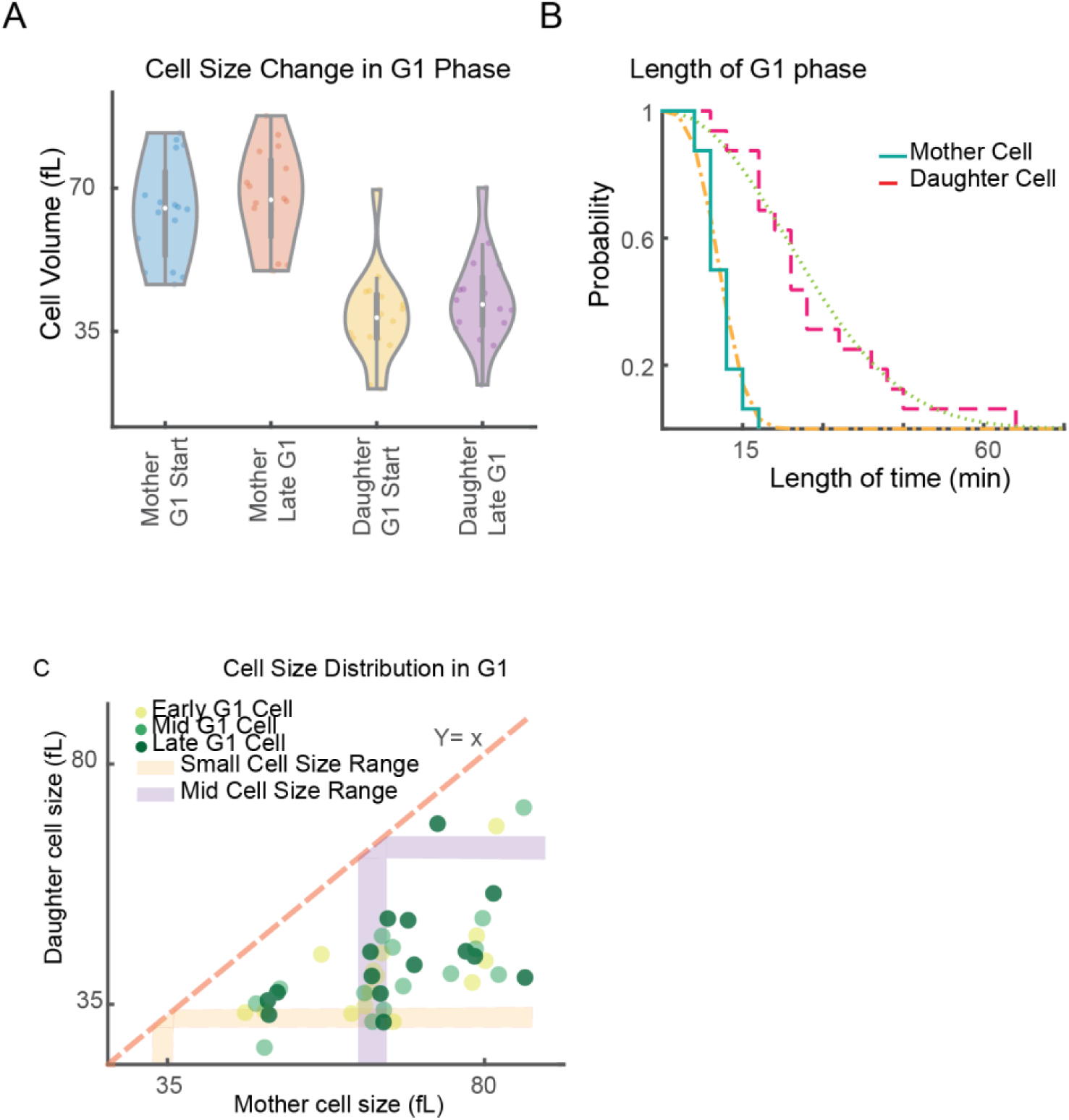
Comparison of cell volume and G1 duration between mother and daughter cells. (A) Cell size distributions at G1 entry and late G1 for Whi5-mCitrine in mother (blue, red) and daughter (yellow, purple) cells. Daughters exhibit more G1 growth. (B) Survival probability of cells remaining in G1 as a function of time. Daughters spend longer in G1 than mothers. (C) Scatter plot of initial vs final cell volume in G1. G1 growth is poorly correlated with starting size in both mothers and daughters.

**Figure S5.**
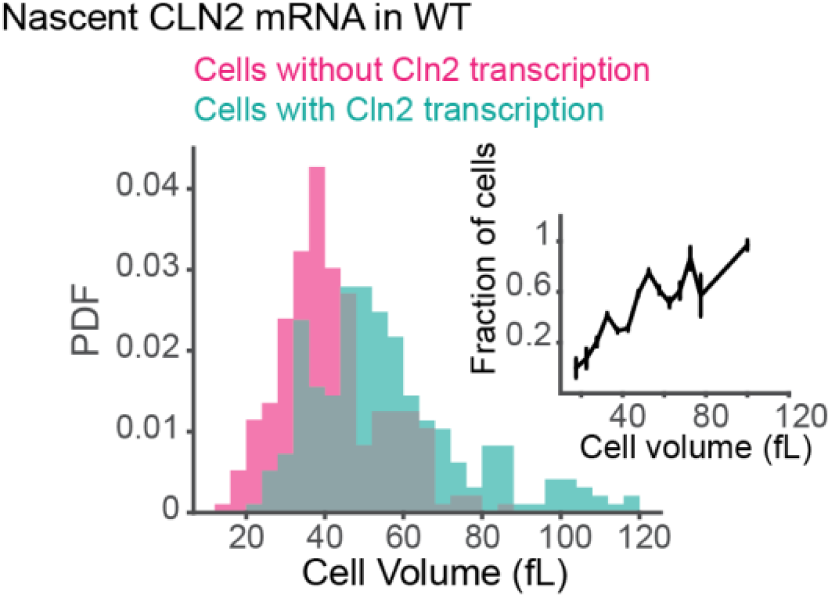
Cell-size dependence of nascent CLN2 transcription frequency. (A) Distribution of cell volumes in which CLN2 nascent mRNA was detected (blue) or absent (pink) by smFISH in WT cells (n = 504). (B) Inset: Fraction of cells with active CLN2 transcription as a function of cell volume, indicating a size-dependent activation pattern.

**Figure S6.**
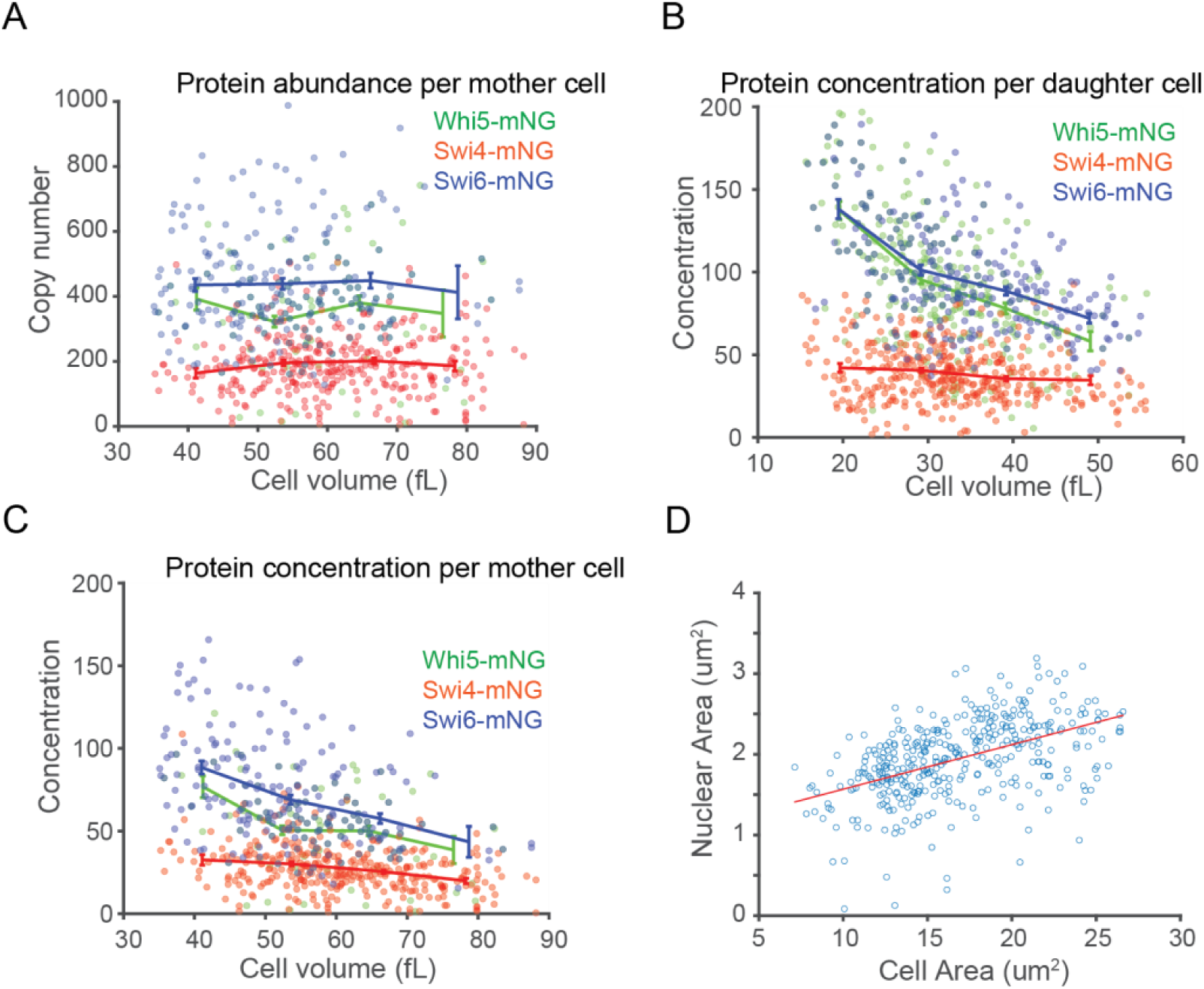
Estimation of protein abundance and concentration as a function of cell volume. (A) Protein abundance (molecule count) per mother cell for Whi5-mNG (n = 171), Swi4-mNG (n = 229), and Swi6-mNG (n = 177) plotted against cell volume. (B) Protein concentration (molecules/fL) per daughter cell as a function of cell volume. Nuclear volume estimated for concentration calculation. (C) Protein concentration per mother cell for the same strains as in (B). (D) Correlation between nuclear and cell volume. The fitted linear equation is y = 0.055 + 1.02x with R^2^ = 0.2344. The average nuclear-to-cell volume ratio is 0.1197.

**Figure S7.**
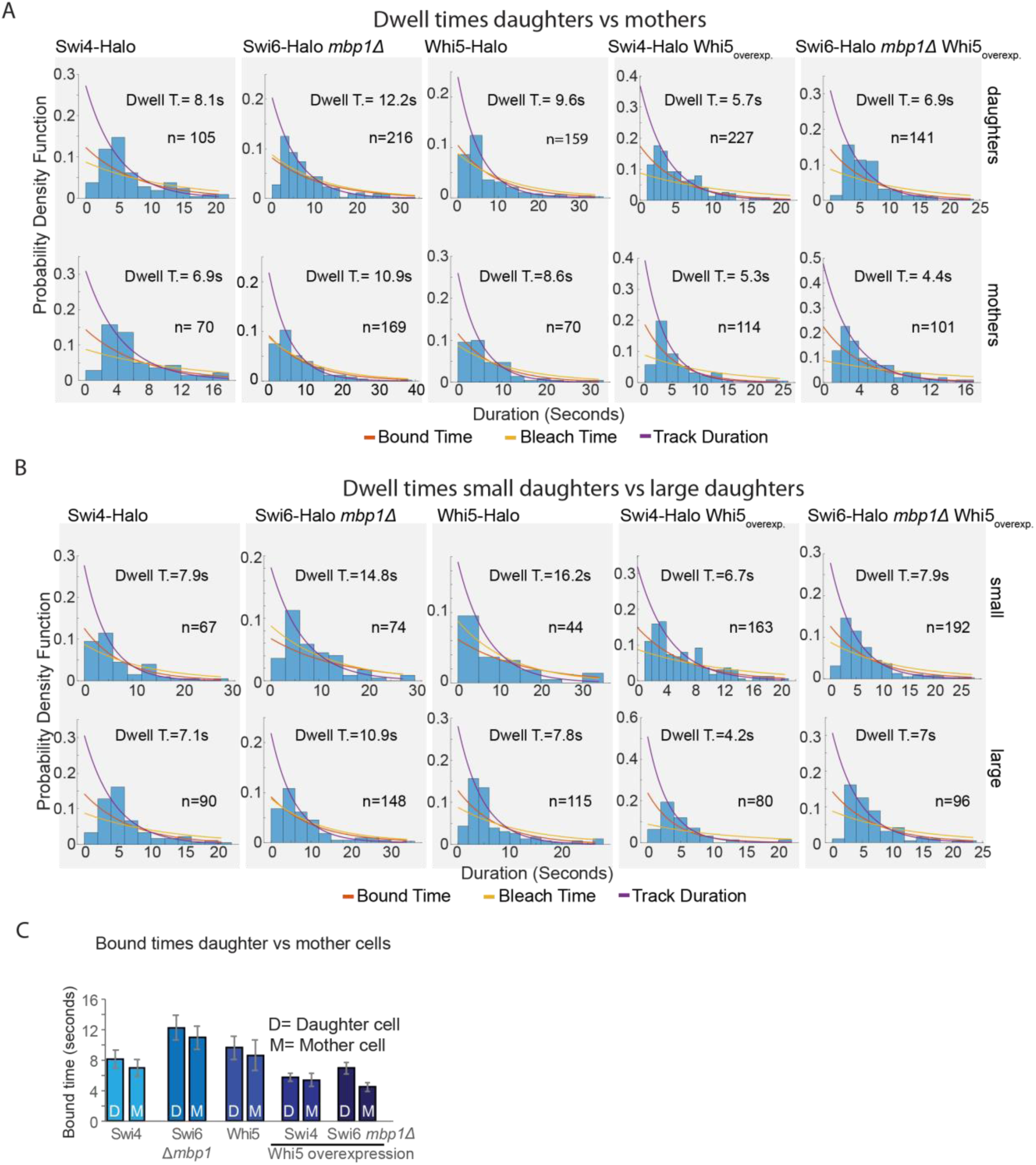
Compare chromatin dwell-time distributions. (A) Chromatin-bound dwell-time PDFs for G1 daughter (top) and mother (bottom) cells. Genotypes: Swi4-Halo, Swi6-Halo mbplΔ, Whi5-Halo, Swi4-Halo with Whi5 overe×pression, and Swi6-Halo mbplΔ with Whi5 overexpression. Superimposed fits: bound-time (red), bleach-time (orange), total track duration (purple). Mean dwell time (T) and sample size (n) shown. (B) Same as in A, but comparing small (<30 fL) versus large (>45 fL) G1 daughter cells. All distributions were normalized and fit with a global two-state exponential decay model accounting for photobleaching. (C) Summary of dwell times comparing daughter against mother cells, data obtained from distributions shown in B. Error bars represent the standard error of the mean.

**Figure S8.**
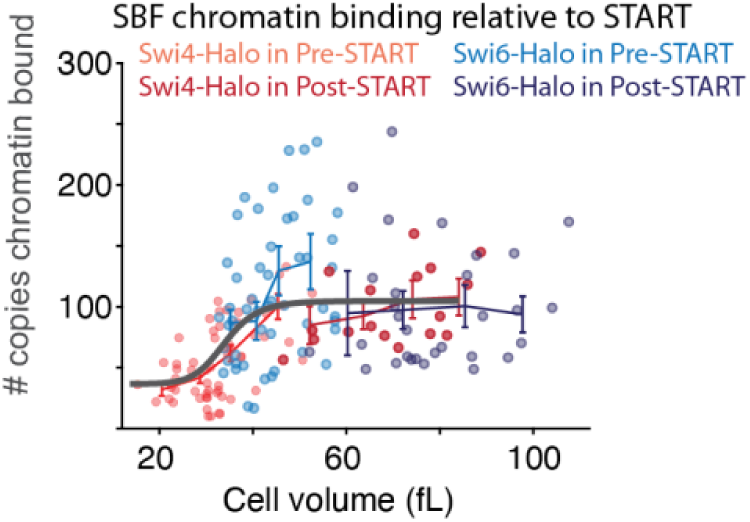
SBF chromatin occupancy shows switch-like behavior at Start. The number of chromatin-bound copies of SBF components Swi4-Halo (red) and Swi6-Halo (blue) is plotted against total cell volume for individual daughter cells classified as pre-START (lighter symbols) or post-START (darker symbols) using Whi5 nuclear export as the benchmark. Solid curves: moviπg-window medians (Loess fit) with 95 % confidence intervals. Error bars on binned means (10 fL bins) illustrate mean ± s.e.m. A sigmoidal increase in SBF occupancy precedes START, followed by a plateau post-START.

**Table S1.**
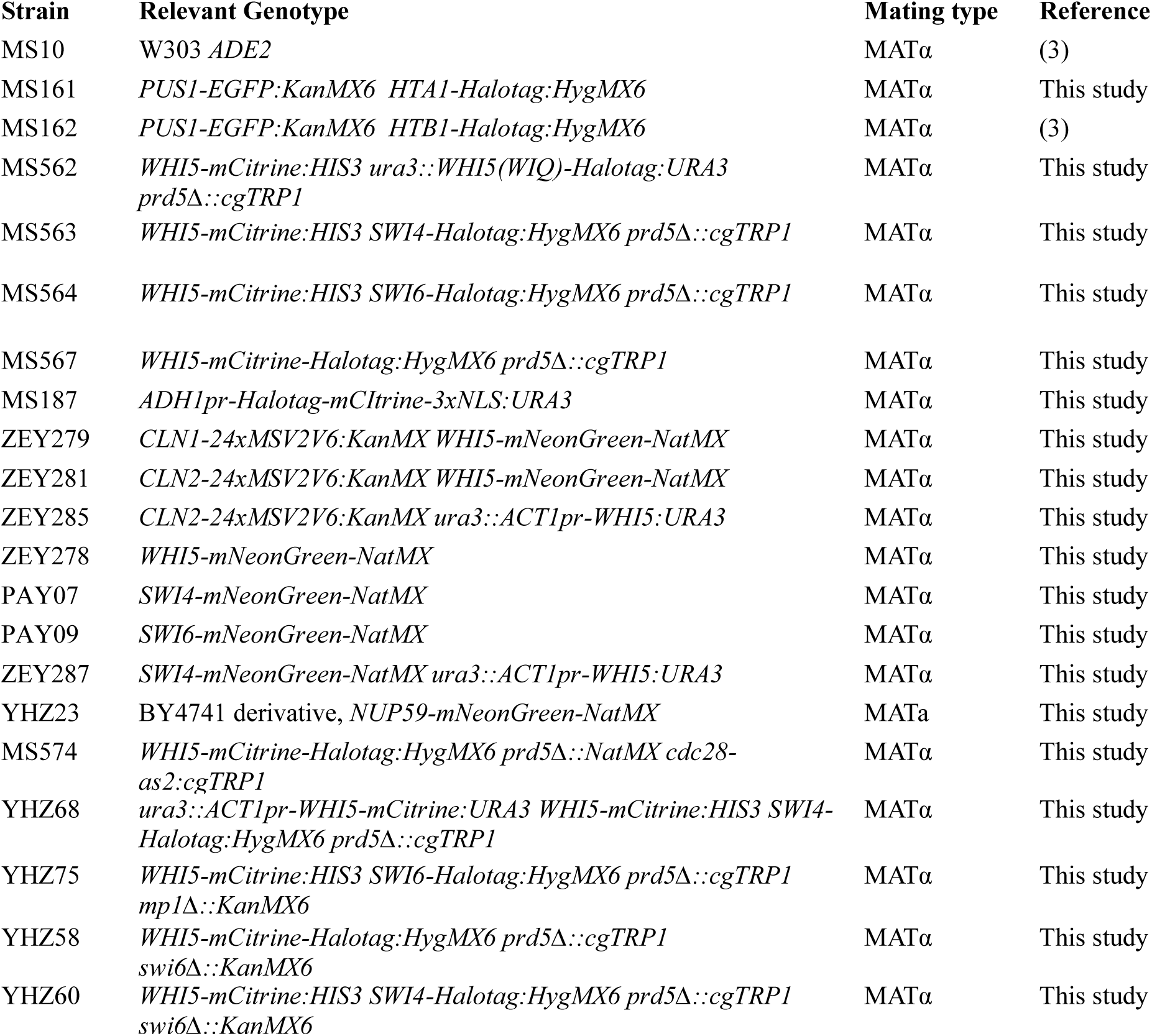
Strains used in this work.

**Table S2.**
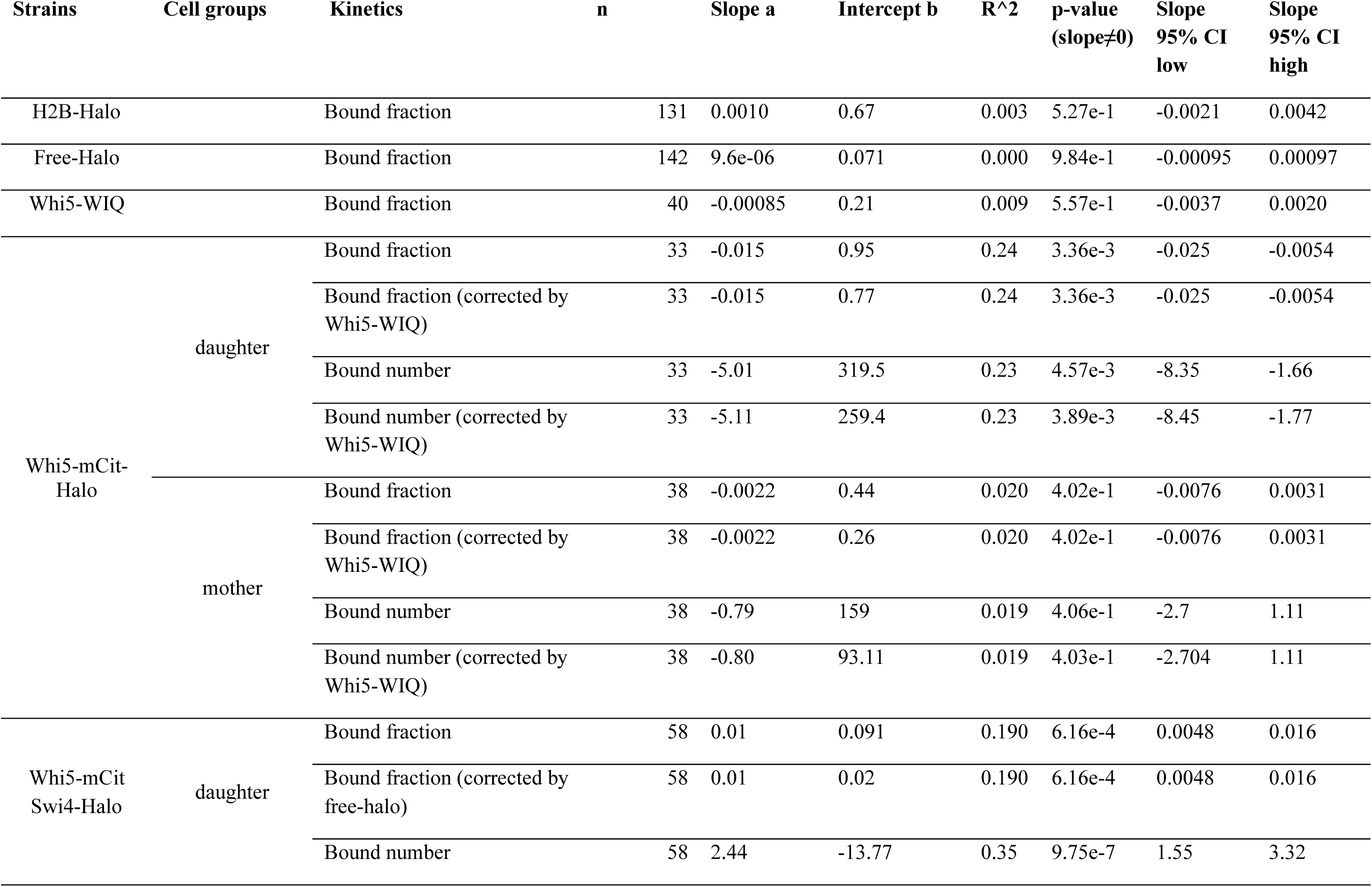
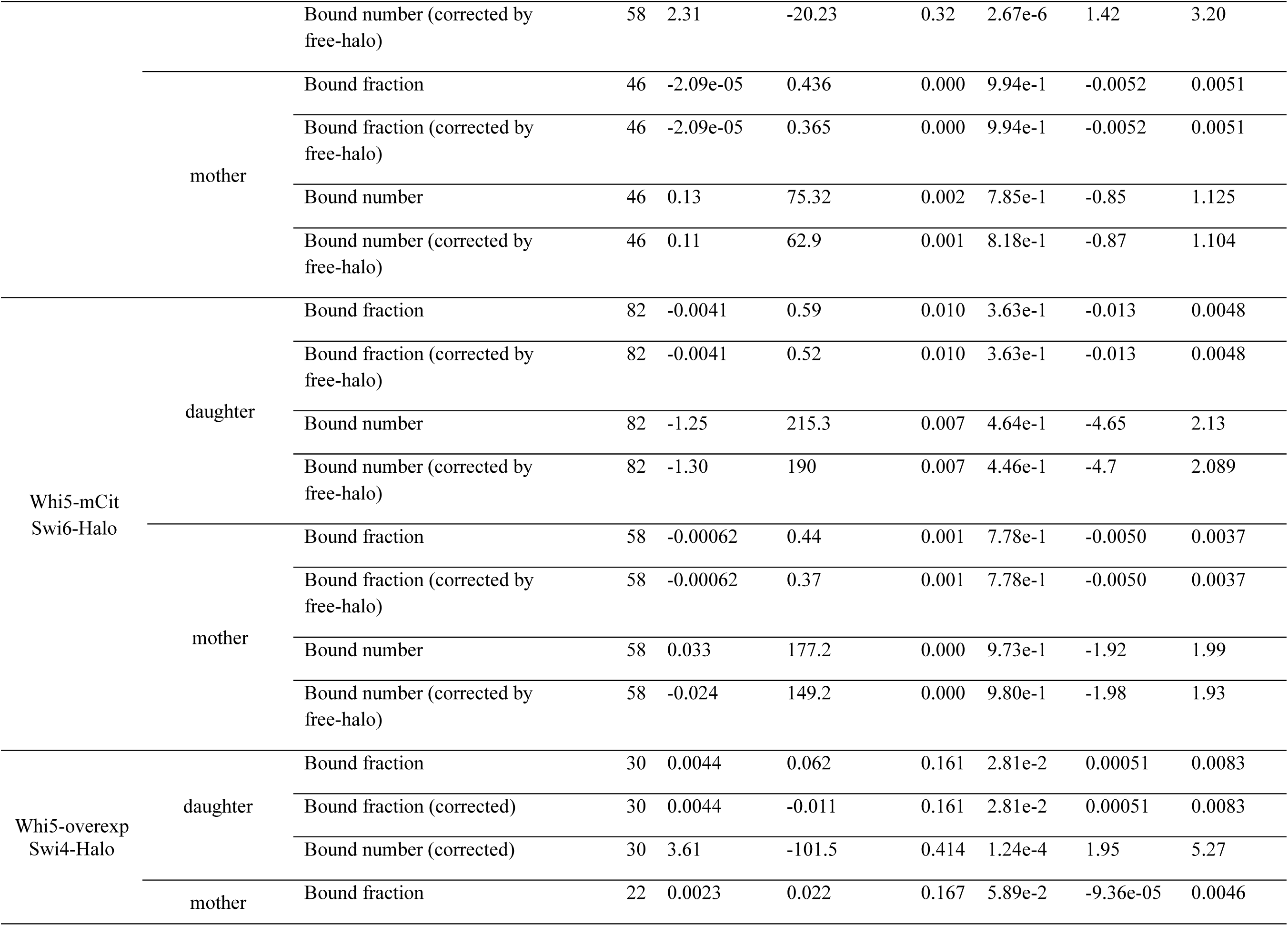
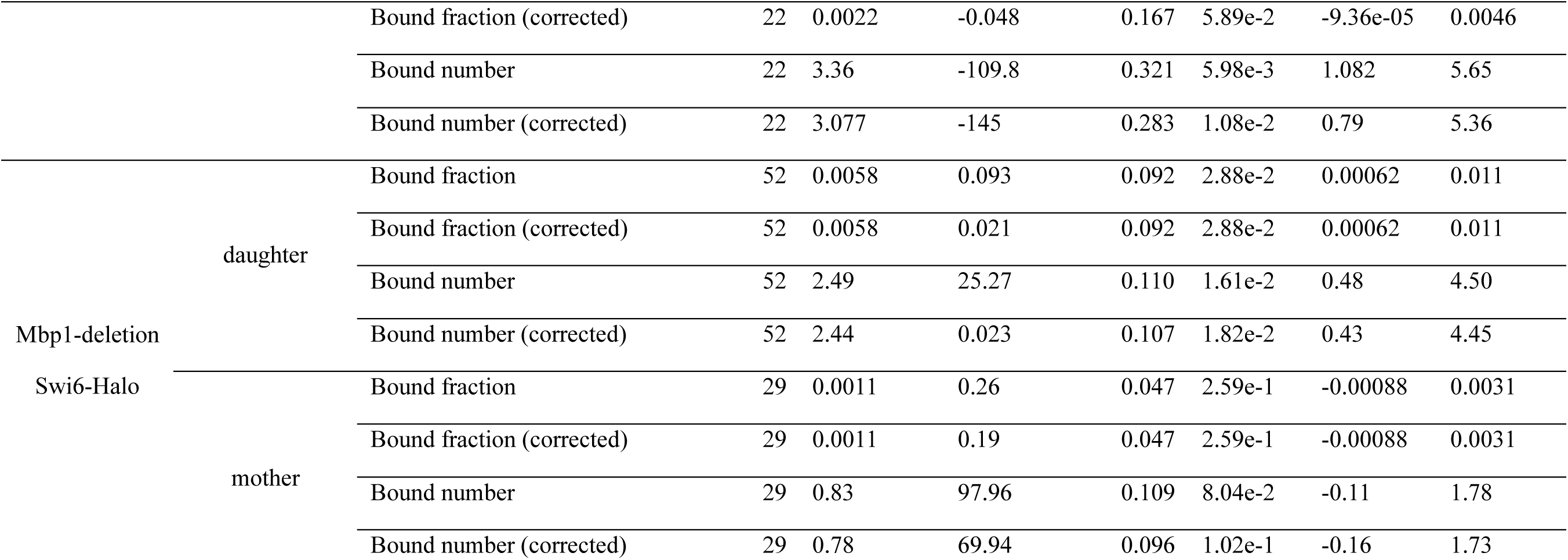
Statistics for linear regression of fast-capture binding fraction versus cell volume. For each strain and cell type (mother or daughter), the indicated response variable *y* (e.g., bound fraction, bound number) was fit as a linear function of cell volume x(fL) using the model *y* = *ax* + *b.* The table reports the number of cells included in each fit (*n*), the best-fit slope *a* and intercept b, the coefficient of determination *(R_2_)* as a measure of goodness of fit, the two-sided p-value testing *a* ≠ *0*, and the 95% confidence interval for the slope.

**Table S3.** Binding kinetic parameters across strains and cell-size groups. Cells were grouped into small or large size bins as indicated. For each strain and size class, *n* denotes the number of cells analyzed in dwell time analysis. Lifetime (s) reports the mean track lifetime. DwellTime (s) is the estimated bound-state residence time; CI_L and CI_H are the lower and upper 95% confidence limits for DwellTime. DwellT_SE is the standard error of the Dwell Time estimate. BoundF is the bound fraction, with SE its standard error. Search Time (s) denotes how long it takes for a molecule to find its targets; Kon is the apparent association rate defined as 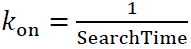 with SearchTime_SE and Kon_SE denoting standard errors.

| Strain name | Size | n | Lifetime | Dwell<br>Time | CI_<br>L | CI_<br>H | DwellT_S<br>E | Bound<br>F | SE | SearchTime<br>e | Kon | SearchTime_S<br>E | Kon_S<br>E |
| --- | --- | --- | --- | --- | --- | --- | --- | --- | --- | --- | --- | --- | --- |
| Whi5-mCit-Halo | Small | 44 | 7.15 | 16.25 | 8.49 | 34.96 | 5.87 | 0.43 | 0.080 | 21.89 | 0.046 | 10.67 | 0.022 |
|  | Large | 115 | 4.85 | 7.81 | 5.50 | 10.73 | 1.33 | 0.12 | 0.039 | 59.55 | 0.017 | 24.82 | 0.007 |
| Whi5-mCit-Swi4-Halo | Small | 67 | 4.90 | 7.95 | 5.26 | 12.30 | 1.72 | 0.32 | 0.028 | 16.81 | 0.059 | 4.23 | 0.015 |
|  | Large | 90 | 4.55 | 7.06 | 5.24 | 9.59 | 1.09 | 0.49 | 0.043 | 7.26 | 0.138 | 1.68 | 0.032 |
| Whi5-mCit-Swi6-Halo | Small | 74 | 6.86 | 14.85 | 9.63 | 22.37 | 3.19 | 0.39 | 0.039 | 22.65 | 0.044 | 6.11 | 0.012 |
|  | Large | 148 | 5.90 | 10.95 | 8.05 | 14.88 | 1.72 | 0.36 | 0.030 | 19.73 | 0.051 | 4.03 | 0.010 |
| Whi5-overexp-Swi4-Halo | Small | 163 | 4.40 | 6.71 | 5.33 | 8.22 | 0.74 | 0.19 | 0.024 | 29.36 | 0.034 | 5.67 | 0.007 |
|  | Large | 80 | 3.16 | 4.20 | 3.03 | 6.03 | 0.74 | 0.25 | 0.023 | 12.74 | 0.079 | 2.74 | 0.017 |
| Whi5-overexp-Swi6-Halo | Small | 192 | 4.89 | 7.91 | 6.30 | 9.76 | 0.88 | 0.24 | 0.023 | 25.46 | 0.039 | 4.31 | 0.007 |
|  | Large | 96 | 4.54 | 7.03 | 5.32 | 9.27 | 0.99 | 0.36 | 0.034 | 12.77 | 0.078 | 2.62 | 0.016 |

**Table S4A.** Proteins copy number linear regression fit.

| Strain name | Cell type | n | a | b | R<sup>2</sup> | p(a>0) | 95% CI |
| --- | --- | --- | --- | --- | --- | --- | --- |
| Whi5-mNeonGreen | daughter | 209 | 2.54 | 22.13 | 0.53 | 1.45e-35 | [2.21, 2.87] |
|  | mother | 171 | 3.29 | -6.69 | 0.77 | 1.89e-56 | [3.02, 3.57] |
| Swi4-mNeonGreen | daughter | 225 | 2.93 | -9.29 | 0.66 | 3.08e-54 | [2.66, 3.21] |
|  | mother | 229 | 2.71 | 1.094 | 0.71 | 8.84e-63 | [2.48, 2.93] |
| Swi6-mNeonGreen | daughter | 197 | 6.98 | 4.67 | 0.85 | 3.01e-82 | [6.56, 7.4] |
|  | mother | 177 | 7.97 | -50.69 | 0.93 | <1e-99 | [7.65, 8.3] |
| Swi4-mNeonGreen (Whi5 overexpression) | daughter | 117 | 7.85 | -25.96 | 0.79 | 6.42e-41 | [7.1, 8.6] |
|  | mother | 111 | 7.42 | -56.67 | 0.77 | 3.08e-37 | [6.66, 8.18] |

**Table S4B.** Proteins concentration linear regression fit. Statistics for linear regression of copy number versus cell volume. Linear model: *y* = *ax* + *b.* Reported are slope *a,* intercept *b,* R*_2_*, and the one-sided p-value testing *a* > 0(or state explicitly if two-sided). 95% CI refers to the slope unless otherwise noted.

| Strain name | Cell type | n | a | b | R<sup>2</sup> | p(a>0) | 95% CI |
| --- | --- | --- | --- | --- | --- | --- | --- |
| Whi5-mNeonGreen | daughter | 209 | -0.022 | 3.99 | 0.089 | 1.00e+0 | [-0.032,<br>-0.013] |
|  | mother | 171 | -0.00077 | 3.19 | 0.001 | 6.30e-1 | [-0.0053,<br>0.0038] |
| Swi4-mNeonGreen | daughter | 225 | 0.013 | 2.21 | 0.038 | 1.62e-3 | [0.0043,<br>0.021] |
|  | mother | 229 | 8.052e-05 | 2.72 | 0.000 | 4.82e-1 | [-0.0034,<br>0.0036] |
| Swi6-mNeonGreen | daughter | 197 | -0.0066 | 7.40 | 0.009 | 9.03e-1 | [-0.017,<br>0.0034] |
|  | mother | 177 | 0.0064 | 6.66 | 0.048 | 1.63e-3 | [0.0022,<br>0.011] |
| Swi4-mNeonGreen (Whi5 overexpression) | daughter | 117 | 0.0065 | 6.98 | 0.006 | 1.97e-1 | [-0.0086,<br>0.022] |
|  | mother | 111 | 0.0056 | 6.25 | 0.019 | 7.23e-2 | [-0.0020,<br>0.013] |

